# Invasion implies substitution in ecological communities with class-structured populations

**DOI:** 10.1101/773580

**Authors:** Tadeas Priklopil, Laurent Lehmann

## Abstract

Long-term evolution of quantitative traits is classically and usefully described as the directional change in phenotype due to the recurrent fixation of new mutations. A formal justification for such continual evolution ultimately relies on the “invasion implies substitution”-principle. Here, whenever a mutant allele causing a small phenotypic change can successfully invade a population, the ancestral (or wild-type) allele will be replaced, whereby fostering gradual phenotypic change if the process is repeated. It has been argued that this principle holds in a broad range of situations, including spatially and demographically structured populations experiencing frequency and density dependent selection under demographic and environmental fluctuations. However, prior studies have not been able to account for all aspects of population structure, leaving unsettled the conditions under which the “invasion implies substitution”-principle really holds. In this paper, we start by laying out a program to explore and clarify the generality of the “invasion implies substitution”-principle. Particular focus is given on finding an explicit and functionally constant representation of the selection gradient on a quantitative trait. Using geometric singular perturbation methods, we then show that the “invasion implies substitution”-principle generalizes to well-mixed and scalar-valued polymorphic multispecies ecological communities that are structured into finitely many demographic (or physiological) classes. The selection gradient is shown to be constant over the evolutionary timescale and that it depends only on the resident phenotype, individual growth-rates, population steady states and reproductive values, all of which are calculated from the resident dynamics. Our work contributes to the theoretical foundations of evolutionary ecology.

## 1 Introduction

A central theme in evolutionary biology is to understand how organisms have evolved to become adapted to their environment. Of particular relevance is to understand adaptation to biotic environments which contain, and are altered by, the interactions of the organism with members of its own and other species (Pásztor et al., 2016; Estrela et al., 2018). Examples of such interactions permeate the biological world but are likely to lead to complex frequency- and density-dependent evolutionary dynamics. It may thus be felt that in general not much can be said about the evolutionary adaptive trajectory of traits.

Notwithstanding this complexity, it is generally argued on theoretical grounds that when mutations cause only small changes to the phenotype under selection, evolutionary change is continual, proceeding by a gradual, small-step by small-step transformation of the phenotype under focus (e.g., Hamilton, 1964; Eshel, 1983; Metz et al., 1995; Rousset, 2004). Such a paradigmatic Darwinian process (e.g., Dawkins, 1986, 1997) relies on the recurrent application of the “invasion implies substitution”-principle, which is the ultimate fixation in the population of any mutant allele being favored by selection when initially rare (i.e., a selective sweep obtains). It has been suggested that the “invasion implies substitution”-principle holds generally (Rousset, 2004; Durinx et al., 2008; Metz and de Kovel, 2013; Lehmann and Rousset, 2014) and has been called a “gift from God” (Hamilton, 1988). The current literature, however, has not been able to account for all aspects of population structure thus leaving unsettled the general validity of the “invasion implies substitution”-principle.

In this paper, we start by introducing and summarising the main theoretical ideas underlying the “invasion implies substitution”-principle (as this has been considered in different literatures and using different approaches) and outline a program that aims at formalizing the “invasion implies substitution”-principle in spatially and demographically structured populations with a specific focus on finding a representation for a selection gradient. We then contribute to this program by proving the “invasion implies substitution”-principle for a scalar-valued quantitative trait in haploid demographically class-structured population that is part of a larger well-mixed ecological community. In so doing, we lay out in detail the concepts of singular perturbation theory and multiple timescale analysis (Fenichel, 1979; Kuehn, 2015).

The intuitive argument for justifying “invasion implies substitution”-principle relies on considering two alleles, a wild-type (resident) allele coding for some phenotype and a mutant allele coding for some closely similar phenotype. The argument is that similar phenotypes lead to (nearly) equally strong interactions between individuals, irrespective of their phenotype, and hence to weak selection. Consequently, the dynamics of a mutant allele frequency *p* in the population is much slower than the dynamics of all other variables governing the demographic and genetic make-up of the population, such us population densities, distribution of demographic classes and genetic associations like relatedness or linkage disequilibria (see Figure 1 panels A and B and Rousset, 2004, p. 196 and p. 206-207 for an early general discussion of this argument). The genetic and ecological variables that operate in fast population dynamical time (collectively referred to as population dynamical variables) can therefore be assumed constant at the slow micro-evolutionary time at which the mutant frequency *p* changes. More precisely, the expected change Δ*p* in mutant frequency *p* is supposed to follow a dynamical equation like

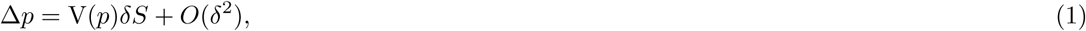

where V(*p*) is a frequency-dependent but always positive measure of genetic variation at the loci under selection. Moreover, *δ* characterises the phenotypic deviation between mutant and resident phenotype and *S* is a frequency-independent selection gradient. Whenever *S* is non-zero, (1) says that if mutant frequency *p* increases when rare as a result of selection, it substitutes the resident; that is, it substitutes its ancestral phenotype. This is the “invasion implies substitution”-principle.

**Figure 1:**
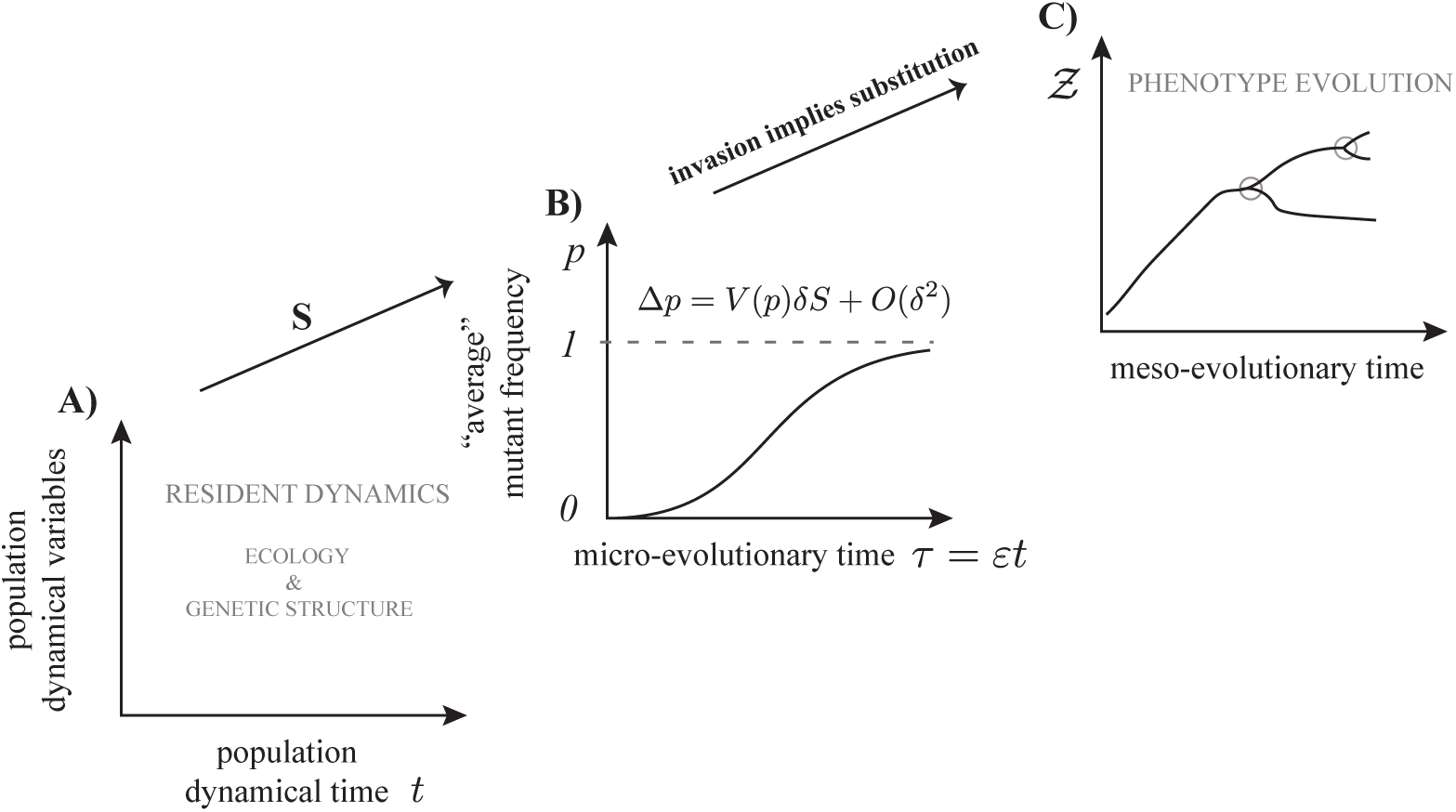
The three timescales that are relevant for the “invasion implies substitution”-principle. A) The population dynamical timescale at which all fast population dynamical variables converge to their steady state. B) The micro-evolutionary timescale at which the weighted average mutant frequency *p* changes, and where the mutant allele may or may not substitute its ancestral resident allele. Whenever the selection gradient *S* is positive, invasion of a mutant will imply substitution (as depicted in the panel). *S* is evaluated at the steady state of the fast population dynamical variables. C) The meso-evolutionary timescale at which the phenotype under selection changes (Metz, 2011). The trait under selection takes values in the trait space 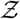. This panel gives the timescale of the trait substitution sequence where each individual trait substitution is defined as an invasion implies substitution event. The “invasion implies substitution”-principle holds whenever *S* is non-zero (gray circles indicate where *S* is zero).

In structured populations, however, finding the required slow dynamical variable *p* is not straightforward. For example, it is known that when individuals are structured into discrete demographic classes such as different age or size classes or spatial locations, the (arithmetic) mean mutant frequency in the population is not a purely slow dynamical variable (Leturque and Rousset, 2002; Rousset and Ronce, 2004) and thus changes across timescales. Moreover, when individuals are structured into continuous or countably infinite demographic classes or spatial locations, population dynamical variables such as population densities or genetic associations need not be fast either (Rousset, 2006; Gyllenberg, 2007, but see Greiner et al. 1994; Cantrell et al. 2017). In such situations a standard timescale separation argument in proving the “invasion implies substitution”-principle is not readily applicable, or, may not even be possible.

Despite of these complications, the timescale separation argument and the “invasion implies substitution”-principle should nevertheless hold in structured populations. This should especially be true not only for arbitrary interactions between individuals, but in cases where populations are spatially and demographically structured into discrete classes, as well as in the presence of environmental fluctuations (Rousset, 2004; Lehmann and Rousset, 2014), when *p* is defined as an average mutant frequency weighted with class-specific reproductive values (e.g., Stubblefield and Seger, 1990; Taylor, 1990). For such population structure the weighted mutant frequency *p* is indeed a purely slow variable (Leturque and Rousset, 2002; Rousset and Ronce, 2004; Rousset, 2004; Lehmann and Rousset, 2014; Grafen, 2015), suggesting that for discrete class-structured populations the dynamics of *p* can be cast in the form (1) and moreover with a selection gradient of the generic form

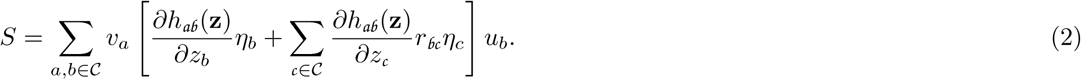

Here, 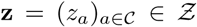 is a class-specific resident phenotype, ***η*** = (*η*_*a*_)_*𝒶*∈𝒞_ gives the direction of the deviation between a mutant and a resident phenotype, and where 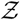 is the phenotype space and 𝒞 is the space of all classes containing a complete description of spatial, demographic, and environmental states an individual can be in (see Figure 2 for the partition of *S* and Section 5 for further discussion). The class-specific growth-rate *h*_*𝒶𝒷*_ (**z**) is an element of a resident growth-rate matrix **H**(**z**) and gives the rate at which a single resident individual in class *𝒷* ∈ 𝒞 that expresses a phenotype *z*_*b*_ produces individuals *𝒶* ∈ 𝒞. The matrix **H**(**z**) has **v** = (*v*_*a*_)_*a*∈𝒞_ and **u** = (*u*_*a*_)_*a*∈𝒞_ as leading left and right eigenvectors with elements giving, respectively, the resident individual reproductive values and the frequency (or probability distribution) of classes, i.e., the class-frequencies. The first directional derivative (*∂h*_*𝒶𝒷*_ (**z**)*/∂z*_*b*_)*η*_*𝒷*_ is taken with respect to the phenotype (more precisely, with respect to the contribution of an allele on the phenotype) of the focal individual whose growth-rate we are considering in *h*_*𝒶𝒷*_ (**z**), and where the weight *η*_*b*_ gives the direction of the deviation between mutant and resident phenotypes as expressed in class *𝒷* ∈ 𝒞. The second directional derivative (*∂h*_*𝒶𝒷*_ (**z**)*/∂z*_*𝒸*_)*η*_*𝒸*_ is taken with respect to the phenotype of individuals in class *𝒸* ∈ 𝒞. These directional derivatives are usually interpreted as fitness effects caused by mutations, and *r*_*𝒷𝒸*_ weights these effects by the genealogical relationship between the focal individual *𝒷* ∈ 𝒞 and an average individual in class *𝒸* ∈ 𝒞. That is, the elements of *r*_*𝒷𝒸*_ are neutral relatedness coefficients (Rousset, 2004). All above quantities are evaluated at the population dynamical steady state. The selection gradient *S* can thus be interpreted as the expected marginal effect (the change in the rate of producing offspring) of an average carrier of the mutant allele.

**Figure 2:**
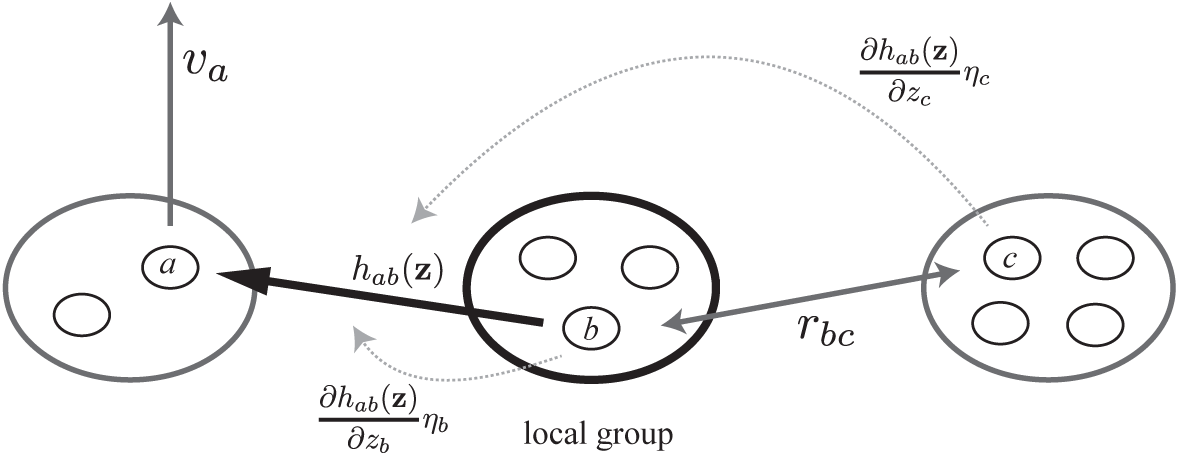
The partitioning of the selection gradient *S* as an inclusive fitness effect. Suppose, as a thought experiment, that when a “button” is pressed all mutant individuals in the population “switch” to expressing the small *δ* deviation. Pressing the button will marginally affect the production rate *h*_*𝒶𝒷*_(**z**) of demographic class *𝒶* offspring by a single mutant individual *𝒷* inhabiting a given discrete spatial location (denoted as “local group”) in two ways. First, it will affect the production “directly” because the mutant individual in class *𝒷* expresses the *δ* deviation, which results in marginal effect 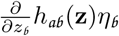. Second, the production will also be affected “indirectly” due to social (frequency or density dependent) interactions with other mutant individuals expressing the *δ* deviation, which results in marginal effect 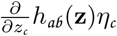 multiplied by the probability that individuals *𝒸* have the mutation (which is conditional on the individual *𝒷* in the local group having the mutation), i.e., the probability *r*_*𝒷𝒸*_. Because *u*_*𝒷*_ is the probability that the individual of interest is in class *𝒷* and the number of future offspring of a single newly born individual *𝒶* is the reproductive value *v*_*𝒶*_, the selection gradient in (2) is obtained from 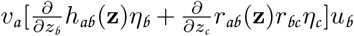 by summing over all possible classes. The selection gradient is thus the marginal effect on mutant allele transmission of a random carrier of the mutant allele having phenotype **z** + *δ****η***.

The “invasion implies substitution”-principle is well established, including several different biological scenarios, for demographically unstructured populations. In particular, it has been shown to hold for haploid well-mixed population models with fluctuating demography (Geritz, 2005; Meszéna et al., 2005; Dercole and Rinaldi, 2008; Dercole and Geritz, 2016), and under limited dispersal in group-structured diploid populations in the absence of demographic fluctuations (Roze and Rousset, 2003; Rousset, 2004, 2006). In all these cases, the selection gradient takes a representation that is a special case of (2). Nevertheless, to date, no detailed proof exists for structured populations specifying all steps leading to (1)-(2) (or some analogue). First and most recently, Lion (2018a,b) discusses the evolutionary dynamics of the trait mean for a polymorphic trait that is tightly clustered around its mean in a demographically class-structured but spatially well-mixed population. Because the timescale separation arguments were made in terms of aggregate variables such as trait means and variances, no complete proof of the “invasion implies substitution”-principle was given that should instead consider the full distribution of mutant frequencies. Second, the “invasion implies substitution”-principle has been considered in the island model of dispersal with finite but demographically fluctuating local population sizes (Rousset, 2004; Rousset and Ronce, 2004; Lehmann et al., 2016), sex-classes with different ploidy levels (Roze and Rousset, 2004) and sex-specific imprinting (Van Cleve et al., 2010). However, no explicit step-by-step proof of the “invasion implies substitution”-principle has been detailed beyond invoking that in these discrete-time models the diffusion approximation method for two timescales (Ethier and Nagylaki, 1980, 1988) applies to them (e.g., Rousset, 2004, p. 196) and thus remains wanting in the literature. In particular, because in demographically structured models the selection gradient can be density-dependent, small perturbations caused by the mutation may lead, e.g. to a catastrophic extinction of the population (Ferriere, 2000; Gyllenberg and Parvinen, 2001; Parvinen, 2016), thus requiring a more detailed analysis on the robustness of the evolutionary mutant frequency dynamics under small but non-zero perturbations caused by the invasion of the mutant. Finally, recent work by Cantrell et al. (2017, Sections 5 and 9.3) discusses the “invasion implies substitution”-principle in populations that are structured along a spatial continuum under local and non-local dispersal. However, they do not provide an explicit expressions for the change in average mutant frequency (1) nor a representation for the selection gradient (2).

In summary, no definitive answer has yet been given as to whether the “invasion implies substitution”-principle holds for all biologically relevant scenarios in structured populations. We thus propose the following “invasion implies substitution”-principle program to explore and clarify the adaptive dynamics of closely similar phenotypes. (i) What is the validity and generality of the “invasion implies substitution”-principle in structured populations with respect to the trait space 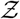 and the class space 𝒞? (ii) If the principle holds in a given model, what conditions must the resident growth-rate matrix (or operator) **H** satisfy, can the micro-evolutionary dynamics of the mutant phenotype systematically be expressed as in (1)? (iii) If the mutant dynamics satisfy (1), can we find an explicit expression for the selection gradient *S* as in (2) (and replacing sums by integrals for continuous trait or class spaces), so that it generically depends on the four aforementioned quantities, **v, u, r**, and **H**?

Our aim in this paper is to contribute to this program and the remainder of it is organised as follows. We start Section 2 by constructing a continuous-time population model that completely describes the population and micro-evolutionary dynamics of the ecological community that is spatially well-mixed but demographically class-structured with a scalar-valued phenotype under selection. Because we will focus on two timescales – a population dynamical and a micro-evolutionary timescale – we will henceforth omit the prefix ‘micro’ unless mentioned otherwise. In Section 3 we move on to study the mutant-resident dynamics in situations where the mutant and its ancestral resident phenotype are closely similar. In Section 4 we proceed to prove the “invasion implies substitution”-principle by decoupling the slow evolutionary dynamics given by mutant frequency weighted by class reproductive values *p* from the fast dynamics given by the population dynamical variables. We conclude by discussing related work and the overall relevance of our results to evolutionary ecology (Section 5).

## 2 Model

Consider an infinitely large haploid well-mixed population where individuals are structured into finitely many demographic classes (Taylor, 1990; Charleworth, 1994), e.g. age or size classes. Each individual is characterized by a single one-dimensional (scalar-valued) continuous trait, which is assumed fixed during its lifespan. The population of interest may also be part of a greater ecological community - individuals of the population interact with individuals from other species (e.g. predator-prey community), which may also be structured into different phenotypes and demographic classes.

### 2.1 Preliminaries

Let 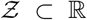 denote the space of phenotypes, 𝒟 the set of distinct demographic classes (which has cardinality *d*), and take time to be continuous. Moreover, suppose that the population is polymorphic with respect to the trait under focus with all in all *k* distinct alleles segregating (each coding for a distinct phenotype), all of which define the resident population. However, because we assume that one (and only one) of the *k* alleles undergoes a mutation giving rise to a new phenotype denoted 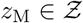 (M stands for mutant), we single out its ancestral phenotype and call it the ancestral resident phenotype 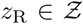, or simply, the resident. After mutation, the population thus consists of a mutant allele (with phenotype *z*_M_), a resident allele (with phenotype 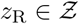), as well as *k* − 1 other alleles, each with their respective phenotypes. Since, under our assumptions, there is a one to one relationship between allele and phenotype, we will generally just speak of mutant and resident phenotypes.

It will be useful to distinguish individuals not only by their phenotype but also the class they are in. For example, a mutant that is in class *𝒶* ∈ 𝒟 will be identified with *z*_M,*𝒶*_. We emphasise that *z*_R,*𝒶*_ and *z*_M,*𝒶*_ take phenotypic value 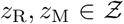, respectively, for all *𝒶* ∈ 𝒟, and that this notation is introduced (only) for a bookkeeping purpose, that is, to keep track of individuals moving in time through the individual-level class space 𝒟. As a consequence, in our model, ***η*** that appears in (2) is just a scalar with a value 1. Finally, to make a distinction between (resident individuals in) resident dynamics and (resident individuals in) mutant-resident dynamics, we will drop out the subscript denoting residents (R) whenever we are discussing ecological communities where the mutant phenotypes are absent.

We now first present a model for a polymorphic resident ecological community where the mutant phenotype is assumed absent (Section 2.2). Then, we extend the model to a situation where one of the phenotypes has undergone a mutation resulting in an arbitrary mutant phenotype and express the dynamical system in terms of class-specific mutant frequencies (Section 2.3). Finally, in Section 2.4, we give several consistency relations and properties that relate mutant-resident dynamics to resident dynamics, which will play a central role in deriving the main results of this paper.

### 2.2 Resident dynamics

Let 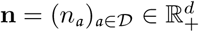 denote the vector of densities (number of individuals per unit space) of (ancestral) resident individuals in all the possible classes the individuals can be in, with element *n*_*𝒶*_ ∈ ℝ_+_ denoting the density of resident individuals in class *𝒶* ∈ 𝒟. Similarly, 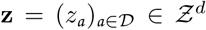 denotes the resident phenotype vector where element *z*_*𝒶*_ identifies individuals in class *𝒶* with a phenotype *z* ∈ 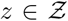. The density vector 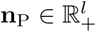 collects, for each class, the density of the other *k* − 1 resident phenotypes in the population of the focal species and the densities of the rest of the ecological community. Hence, if we have a community with a single species *l* = (*k* − 1)*m*, otherwise *l* > (*k* − 1)*m*.

The resident dynamics is given by the set of ordinary differential equations

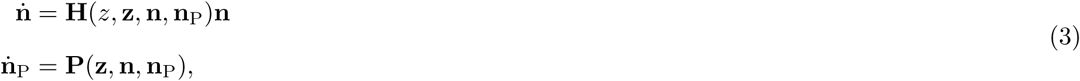

where the dot “·” above the density vectors **n** and **n**_P_ denotes the time derivative “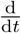”. The matrix **H** = (*h*_*𝒶𝒷*_)_*𝒶,𝒷*∈𝒟_ ∈ ℝ^*d*×*d*^ is the (ancestral) resident growth-rate matrix where entry *h*_*𝒶𝒷*_ (*z*, **z, n, n**_P_) is a sufficiently smooth growth-rate function giving the rate at which a single individual of class *𝒷* ∈ 𝒟 produces individuals of class *𝒶* ∈ 𝒟. We emphasise that the first argument 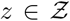 in the growth-rate matrix **H**(*z*, **z, n, n**_P_) identifies the phenotype of the individual whose growth-rate we are considering, while all the remaining arguments describe the environment that the individual finds itself in. The matrix **P** ∈ ℝ^*l*×*l*^ is the growth-rate matrix of the rest of the resident population and the ecological community and is also a function of the environment that the individuals find themselves in. For notational convenience, especially when it is clear from the context, we will drop from the growth-rate matrices and functions all arguments that describe the environment, for example, we may write **H**(*z*) instead of **H**(*z*, **z, n, n**_P_) and **P** instead of **P**(**z, n, n**_P_).

We note that all the growth-rate functions presented in this paper are constructed by assuming an infinitely large well-mixed ecological community, where individuals are assumed to undergo demographic individual-level processes on a Poissonian basis; the demographic processes can be either asocial where individuals react by themselves e.g., dying or moving from one age class to another, or social, resulting from random encounters of pairs of individuals. The probability of any higher order encounter vanishes in continuous-time models. However, all growth-rate functions can be non-linear and of any complexity as we allow for arbitrary frequency and/or density dependent (pairwise) interactions. Different underlying assumptions on the encounters between individuals is possible, facilitating e.g. multiplayer games (Weibull, 1995), but are not dealt with in this paper.

#### 2.2.1 Steady state of the resident dynamics

Throughout the paper we assume that there exists an equilibrium point 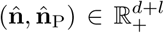 to which the community given by (3) converges to and then stays at. Importantly, this equilibrium is assumed to be hyperbolically stable, i.e. the real part of the dominant eigenvalue of the linearized version of system (3) evaluated at the equilibrium is negative and bounded away from zero (Hirsch et al., 1974). However, we allow the system (3) to contain multiple non-negative equilibria or other attractors at which the community could potentially reside. Assuming multiple equilibria or other attractors is not problematic when considering evolutionary dynamics because the so-called *tube theorem* (Geritz et al., 2002) excludes “attractor switching” for mutant-resident dynamics with closely similar phenotypes. That is, the dynamics of the mutant with a similar phenotype to a resident will never evolve to an alternative attractor. In Section 5 we discuss how our results can be extended to more complicated attractors than equilibria.

### 2.3 Mutant-resident dynamics

We now introduce the mutant phenotype 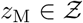 into the resident population. Let 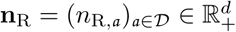 and 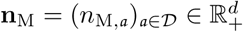 denote the vectors of densities and 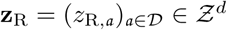 and 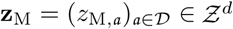 the vectors of phenotypes of (ancestral) residents and mutants, respectively, in all the possible classes the individuals can be in. The mutant-resident dynamics is then given by

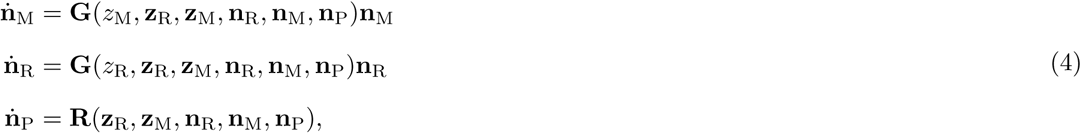

where **G** = (*g*_*𝒶𝒷*_)_*𝒶,𝒷*∈𝒟_ ∈ ℝ ^*d*×*d*^ is the growth-rate matrix of individuals in the mutant-resident population, such that **G**(*x*) := **G**(*x*, **z**_R_, **z**_M_, **n**_R_, **n**_M_, **n**_P_) is the growth-rate matrix of a phenotype *x* ∈ {*z*_M_, *z*_R_} and that each entry *g*_*𝒶𝒷*_ (*x*) is a sufficiently smooth growth-rate function giving the rate at which a single individual with phenotype *x* ∈ {*z*_M_, *z*_R_} in class *𝒷* ∈ 𝒟 produces individuals in class *𝒶* ∈ 𝒟. It is clear from this formulation that as we have assumed the ecological community to be spatially well-mixed, all individuals are surrounded by equal number (density) of mutants and resident and thus the only difference in their growth-rate matrix **G** is due to their own phenotype (the first argument). Similarly to the second line of the resident dynamics (3), **R** ∈ ℝ^*l*×*l*^ is the growth-rate matrix of the *k* − 1 remaining resident phenotypes in each class and of the rest of the ecological community.

#### 2.3.1 Relative mutant-resident dynamics

Because we are interested in the relative dynamics of mutants 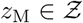 and (ancestral) residents 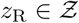, we change the dynamical variables and instead of considering mutant **n**_M_ and resident **n**_R_ densities we consider the frequency of mutants 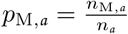 in class *𝒶* ∈ 𝒟, where *n*_*𝒶*_ = *n*_M,*𝒶*_ + *n*_R,*𝒶*_ is the total density of mutants and residents in class *𝒶* ∈ 𝒟. The vectors **p** = (*p*_M,*𝒶*_)_*𝒶*∈𝒟_ ∈ [0, 1]^*d*^ and 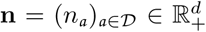, with slight abuse of notation, are thus the vectors for class-specific mutant frequencies and class-specific total densities of (mutant and ancestral resident) individuals, respectively. We emphasise that since we are interested in the relative dynamics of mutants and their ancestral residents, the mutant frequency *p*_M,*𝒶*_ is defined with respect to mutants and their ancestral residents in class *𝒶* ∈ 𝒟, not all *k* resident phenotypes present in the population.

We can now rewrite the mutant-resident dynamics (4) in terms of the class-specific mutant frequencies **p** and the class-specific total population densities **n** as

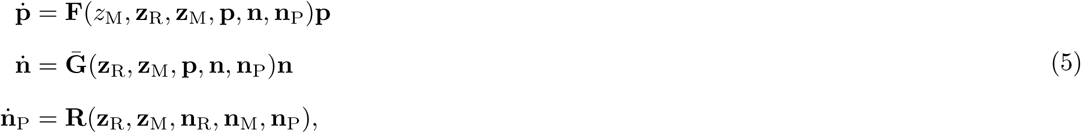

where 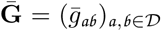, with 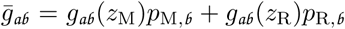, is the average mutant-resident growth-rate matrix, and where **F** = (*f*_*𝒶𝒷*_)_*𝒶,𝒷*∈𝒟_ ∈ ℝ^*d*×*d*^ is the relative growth-rate matrix (see Appendix 6.1 for further details). The entries of the relative growth-rate matrix for mutants **F**(*z*_M_) := **F**(*z*_M_, **z**_R_, **z**_M_, **p, n, n**_P_) are obtained by differentiation

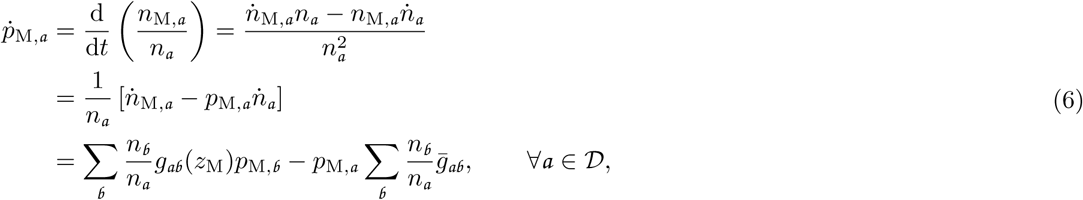

where we have used equations (4) and (5) and the definition of class mutant frequencies *p*_M,*𝒶*_. Motivated by Lion (2018b, Appendix A.3), it will be useful to rewrite (6) by subtracting and adding a term 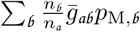, to obtain

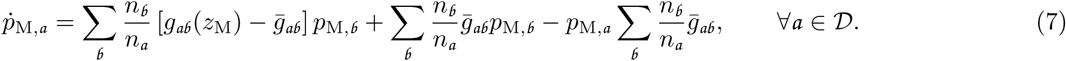

This allows us to partition the mutant relative growth-rate matrix as

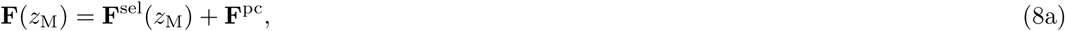

where 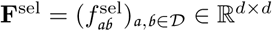 and 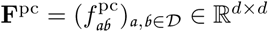 with entries, respectively, given by

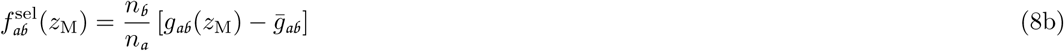

and

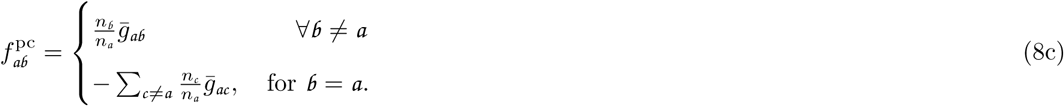

Notice that 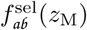 is proportional to the difference between mutant *g*_*𝒶𝒷*_(*z*_M_) and average growth-rates 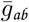 and thus captures the effect of selection (hence the superscript “sel”) on mutant allele frequency change. The second term 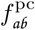 is proportional only to average growth-rates 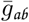 and hence captures non-selective effects on allele frequency change due to transitions between classes. Since the relative growth-rate of an individual due to the term 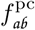 is non-selective and thus independent of ones phenotype (see also Appendix 6.1), the argument present e.g., in 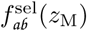 is not included in 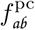, but it should nevertheless be kept in mind that **F**^pc^ depends both on mutant and resident traits. Such non-selective transitions between classes nevertheless affect the dynamics of the mutant frequency, for instance if one class of individuals, say newborns (or individuals living in a good habitat) have higher reproductive success than older individuals (individuals living in bad habitat). Such deterministic change of allele frequency due to non-selective forces have generally been referred to as changes due to “transmission” (following Barton and Turelli, 1991; Kirkpatrick et al., 2002), since they result from alleles changing contexts (e.g., from good habitat to bad habitat, from young to old individual; see Kirkpatrick et al., 2002 for more details on the concept of the context of an allele and a discussion of transmission as an evolutionary force). When the different contexts an allele can reside in are demographic classes, the changes in allele frequency due to transmission have been called “passive changes” (Grafen, 2015; Lion, 2018a,b) and we adhere to this terminology (hence the superscript “pc”).

### 2.4 Properties of growth-rates

In this section we present three properties that relate mutant-resident dynamics (4) to resident dynamics (3) and then we apply them to the mutant relative growth-rate matrix (8). These properties and their applications play a central role in Section 3 when discussing mutant-resident dynamics for closely similar phenotypes and in Section 4 when proving our main result. The consistency relation given below is fully analogous to the relation given in Geritz et al. (2002); Dercole (2016); Dercole and Geritz (2016) and the proposition given below is an analogue to a property derived for unstructured populations (Meszéna et al., 2005; Dercole, 2016, see also Diekmann et al. 2001).

#### Consistency relations

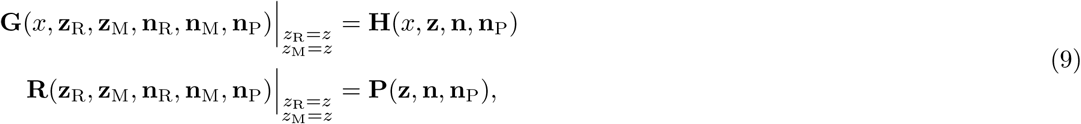

for any 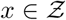. This relation says that the growth-rate of any individual from any population and species in the ecological community, when all (other) individuals in the population are of the same phenotype 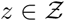, is its growth-rate in a resident ecological community (3) where **n** = **n**_R_ + **n**_M_.

##### Corollary

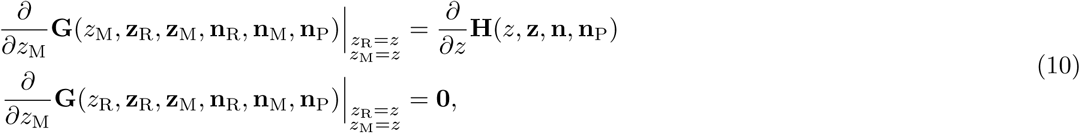

This property follows immediately from the Consistency relation describing the effect that a mutant phenotype of an individual has on its own growth-rate. Trivially, residents do not have a mutant phenotype and so there is no such effect for the resident growth matrix. The same is true also for the matrix **R**, but as we don’t need the Corollary for **R** we haven’t included it here.

##### Proposition

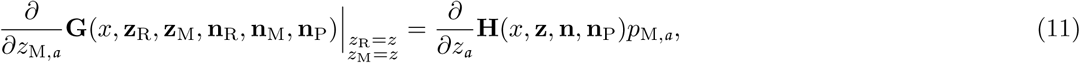

for any 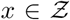 and for all *𝒶* ∈ 𝒟. This property says that the effect that all mutants in class *𝒶* ∈ 𝒟 in the mutant-resident community (4) have on the individual growth-rate (left-hand side of (11)), is equal to the effect that all individuals in class *𝒶* ∈ 𝒟 in the resident community (3) have on the individual growth-rate, weighted with the probability that given a random pairwise encounter with an individual of class *𝒶* ∈ 𝒟, it is a mutant (right-hand side of (11)). This property is a consequence of the growth-rate function being constructed in terms of pairwise interactions between individuals (generalized mass action law), and is a direct generalization of the property 4 given for unstructured populations in Dercole (2016) (see also Meszéna et al., 2005).

#### 2.4.1 Properties of relative growth-rates

Here we apply the above properties (9)-(11) to the mutant relative growth rate matrix (8). Substituting the consistency relation (9) into (8) implies that the selection component of the relative growth-rate matrix **F**^sel^ = **0** is a null matrix for phenotypic equality between mutant and its (ancestral) resident, therefore

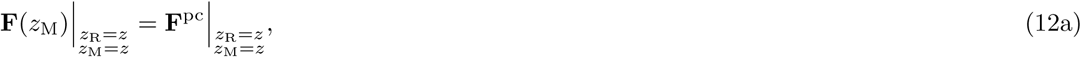

where

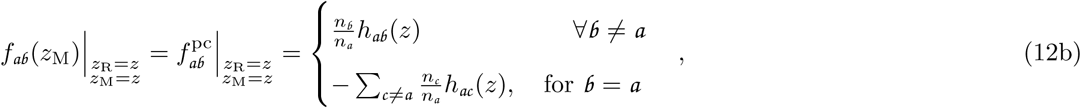

for all *𝒶, 𝒷* ∈ 𝒟. We thus confirm that under phenotypic equality, selection (i.e., the component **F**^sel^(*z*_M_)) plays no role (as it should not) and that the change in class-specific mutant frequencies is non-trivial and purely determined by the matrix **F**^pc^. That is, under phenotypic equality it is the “passive changes” that determines the dynamics of class-specific mutant frequencies (Taylor, 1990; Stubblefield and Seger, 1990; Charleworth, 1994; Grafen, 2015; Lion, 2018a,b).

The Corollary (10) and the Proposition (11) immediately imply, respectively, that

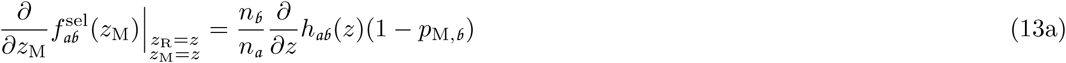

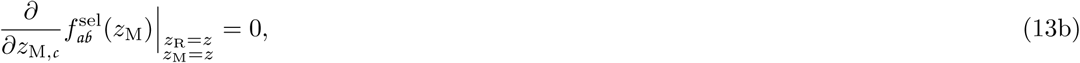

for all *𝒶, 𝒷, 𝒸* ∈ 𝒟. Analogously to above, both properties describe the effect that a mutant phenotype has on the mutant relative growth-rate. The property (13a) follows from the fact that the effect of a mutant phenotype on ones own growth-rate is 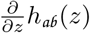 if one is a mutant and 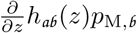 if one is an average (random) individual in class *𝒷* ∈ 𝒟. The property (13b) in turn follows from the fact that in a well-mixed population all individuals experience the exact same social environment and hence the effect that mutants in class *𝒸* ∈ 𝒟 have on a mutant growth-rate and an average growth-rate are equal.

## 3 Mutant-resident dynamics for nearby phenotypes

In this section, we will study the relative mutant-resident dynamics (5) for closely similar phenotypes. To prove the “invasion implies substitution”-principle by using a timescale separation argument, we wish that for closely similar phenotypes the mutant frequency in the population is a much slower dynamical variable than all other dynamical variables in the model. If so, the fast dynamical variables would then have enough time to reach their steady state (or at least to be sufficiently close to it) and thus could be considered as constant arguments of the (much slower) evolutionary dynamics of the mutant frequency. To check the timescale of **p, n, n**_P_ in the relative mutant-resident dynamics (5), let *z*_M_ = *z*_R_ + *δ* and let us Taylor expand (5) up to the first order about *δ* = 0 to obtain

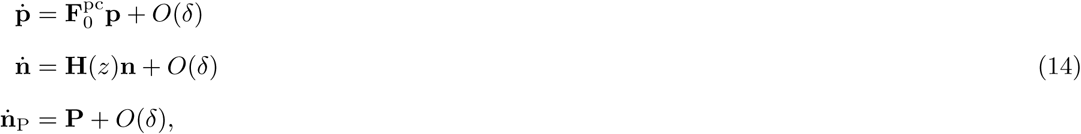

where we have used (9) and (12) and where 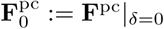 is as given in (12). From these equations, we can see that all variables **p, n** and **n**_P_ have non-zero terms of order *O*(1), and hence are all fast population dynamical variables fluctuating in fast population dynamical time. In other words, none of the dynamical variables **p, n** nor **n**_P_ are (at least not purely) slow dynamical variables dominated by the terms of order *O*(*δ*). This is true in particular for the class-specific mutant frequencies **p** and consequently also for the (arithmetic) mean mutant frequency in the population *p*_M_ = **up** = ∑_*𝒶*_ *u*_*𝒷*_ *p*_M,*𝒶*_, where **u** = (*u*_*𝒶*_)_*𝒶*∈𝒟_ and where 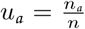 is the frequency of individuals in class *𝒶* ∈ 𝒟 (Appendix 6.2). Since there are no purely slow dynamical variables, a timescale separation can not be readily performed.

In the next Section 3.1, we show that there exists a purely slow dynamical variable and that it is the average mutant frequency weighted by class reproductive values. In the following Section 3.2, we then find the steady state to which the fast population dynamical variables approach to, and then in Section 4 we use these results to prove the “invasion implies substitution”-principle.

### 3.1 The average mutant frequency weighted with class reproductive values

To find a purely slow dynamical variable that tracks changes in class mutant frequencies **p**, thus tracking also the mean mutant frequency in the population *p*_M_, we take an average of **p** over all *p*_M,*𝒶*_ with weights chosen such that the change of this weighted average mutant frequency is zero under phenotypic equality. This will guarantee that under phenotypic similarity the dynamics will be governed by terms of order *O*(*δ*).

To this end, let ***α*** be, for the moment, an arbitrary vector of weights normalized as ∑_*𝒶*∈𝒟_ *α*_*𝒶*_ = 1, and lets denote the average mutant frequency weighted by ***α*** with

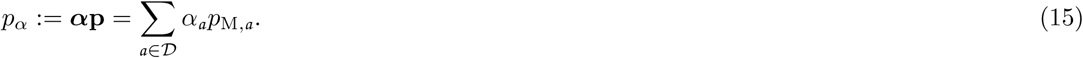

Because we are interested in the dynamics of *p*_*α*_, we differentiate with respect to time *t* and obtain

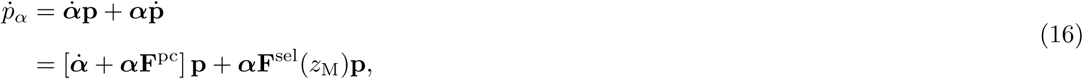

where we have used (5) and (8) (see also for analogues steps taken in Lion 2018b, Appendix A.3). To fulfill the requirement that the change in *p*_*α*_ in (16) is zero under phenotypic equality, we must necessarily have that

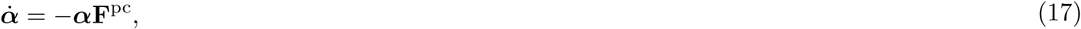

because **F**^sel^|_*δ*=0_ = **0** (Section 2.4.1). Since 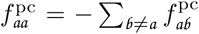, the matrix **F**^pc^ is the transition matrix for a *backward* continuous-time mutant-resident Markov chain ***α*** on the state space 𝒟. Moreover, each element 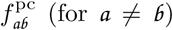 as given in (8) gives the backward in time probability, given a demographic event, that an individual (or its lineage) in class *𝒶* ∈ 𝒟 came from class *𝒷* ∈ 𝒟. This implies that the dynamical equation (17) defines the dynamic version of class reproductive values ***α*** as in the continuous time model of Lion (2018b, p. 624) and the discrete time model of Lehmann (2014, eq. 7).

Using the dynamic definition for class reproductive values (17), the dynamics of the weighted mutant frequency (16) reduces to

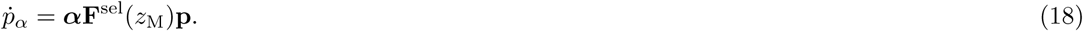

We have thus obtained that since ***α*** by definition satisfies (17), the dynamics of the weighted mutant frequency *p*_*α*_ is determined purely by the selection component of the relative growth rate matrix as given in (18) (see also Appendix 6.5.4). Interestingly, as we have made no assumptions on the magnitude of *δ*, the above equation is valid for arbitrary phenotypic values 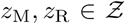 and thus for arbitrary strength of selection. Moreover, because **F**^sel^|_*δ*=0_ = **0** is a null matrix (12), the dynamics of *p*_*α*_ under phenotypic similarity (*δ* small) is

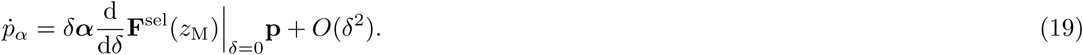

The dynamics of *p*_*α*_ for closely similar phenotypes is thus dominated by the terms *O*(*δ*), and is thus a well suited proxy for the slow evolutionary dynamics of **p** (and *p*_M_).

Finally, we note that the dynamical interpretation (17) is fully consistent with the standard asymptotic definition of class reproductive values that has long been used in class-structured models (e.g., Stubblefield and Seger, 1990; Taylor, 1990; Taylor and Frank, 1996; Leturque and Rousset, 2002; Rousset and Ronce, 2004; Rousset, 2004; Lehmann and Rousset, 2014; Grafen, 2015); at the steady state where (17) is zero, 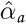 gives the *asymptotic* probability that the ancestral lineage of a random individual was in class *𝒶* ∈ 𝒟. At the steady state in forward in time, 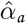 thus gives the fraction of future individuals that descend from individuals in class *𝒶* ∈ 𝒟. Moreover, by defining 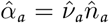, the so-called *individual* reproductive value 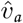 gives the fraction of individuals in distant future that descend from a single individual in class *𝒶* ∈ 𝒟, or with an alternative scaling where 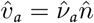 and where 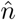 is the total population size, 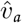 gives the *number* (density) of future offspring that descend from a single individual *𝒶* ∈ 𝒟. Reproductive values, whatever is the scaling, thus define the long-term contribution of individuals to the future composition of the population.

### 3.2 Steady states and the critical and slow manifolds

In Section 3.1, we found that the slow evolutionary dynamics of the weighted average mutant frequency *p*_*α*_ is a slow dynamical variable dominated by the terms of order *O*(*δ*), and that it is a function of the fast population dynamical variables ***α*, p, n** and **n**_P_ governed by the terms of order *O*(1) (Section 3). Under phenotypic equality (*δ* = 0), the dynamics of the ecological community is thus fully described with

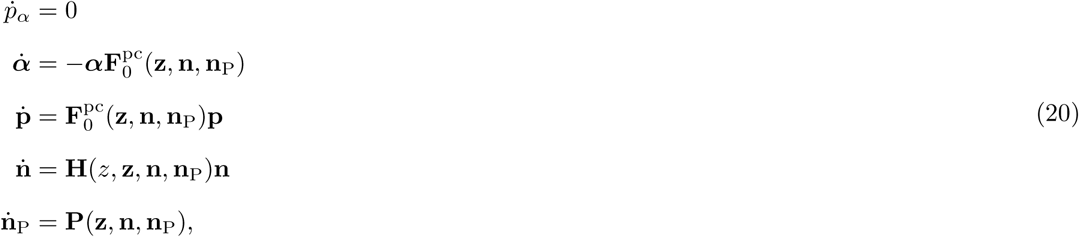

where we used (14), (18) and where we have for clarity included all the arguments. Therefore, in fast population dynamical time, the variables (***α*, p, n, n**_P_) fluctuate and are expected to reach their steady state while the weighted mutant frequency *p*_*α*_ stays constant. The steady state 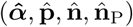 of (20) must, by definition, satisfy

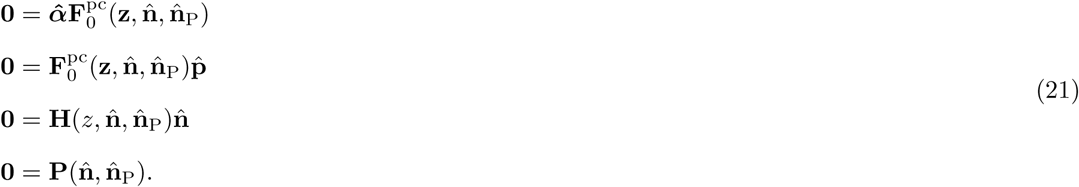

We recall from Section 2.2.1 that the equilibrium solution 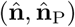 for the bottom two equations exists and is hyperbolically stable (by assumption). Moreover, since the matrix 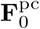 is linear in the elements of **p** (8), the steady states of 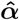 and 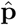 must also exist (see below). The remaining task is to find an explicit expression for the steady state 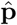 (we do not need one for the rest of the variables), which can be solved from

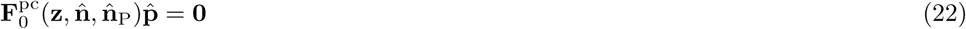

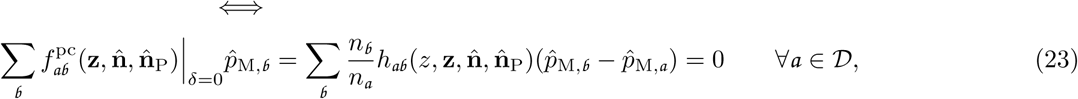

and is given by

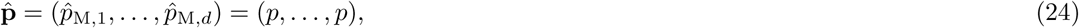

where the class-specific mutant frequencies *p*_M,*𝒶*_ in all classes *𝒶* ∈ 𝒟 are equal. Notice that any value of *p* (biologically meaningful values lie between 0 and 1) gives a solution to (22) and hence the complete solution to (22) consists of infinite number of equilibria (which lie on a line; note that such a degenerate solution results from the fact that 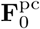 is a non-invertible matrix, Appendix 6.3). The exact value of *p* ∈ [0, 1] to which the class mutant frequencies *p*_M,*𝒶*_ approach to, ∀*𝒶* ∈ 𝒟, depends on the initial condition **p**(*t* = 0). Interestingly, since by definition *p*_*α*_(*t*) = ***α***(*t*)**p**(*t*) for all *t* as given in (15), and since under phenotypic equality the weighted average frequency is constant in fast population dynamical time (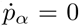 as shown in Section 3.1, but note that in slow time *p*_*α*_ is no longer a constant), we must have that 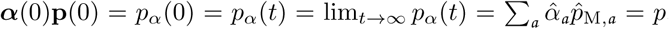 and so 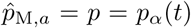 for all *a* ∈ 𝒟 and for all *t*; that is, the asymptotic mutant allele frequency in each class is equivalent to the reproductive value weighted initial mutant frequency.

#### 3.2.1 Critical manifold

Above we obtained that whenever the mutant and resident phenotypes are equal *δ* = 0, the dynamics given by system (20) approaches in fast population dynamical time a steady state 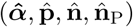 that can be solved from (21). We represent the infinite number of equilibrium points 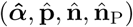 satisfying (21) as the set

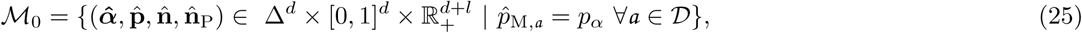

where Δ^*d*^ is the *d*th simplex and the subscript 0 indicates that we are studying the dynamics for the case where *δ* = 0 (see Figure 3, top panel). The set ℳ_0_ is the so-called critical (or equilibrium) manifold (Jones 1995, Definition 1, p. 49; Kuehn 2015, p. 12, see also Appendix 6.4.2 and recall that a manifold is here a sub-space of the original state space). This ℳ_0_ manifold defines in fast population dynamical time the set of equilibrium (or critical) points to which the dynamical system with phenotypic equality *δ* = 0 approaches to. As such, it can be thought of as the state space for the average weighted mutant frequency *p*_*α*_ when *δ* = 0 (see Figure 3). Because 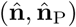 is hyperbolic and the critical manifold ℳ_0_ is compact (the set of points are bounded and closed) consisting of a neutral line of equilibria, it follows that ℳ_0_ is compact and a normally hyperbolic invariant manifold (Appendix 6.4.2). Roughly speaking, an invariant manifold (i.e. a manifold where the dynamical system maps its points onto possibly other points on the manifold) is normally hyperbolic if the dynamics near the manifold is governed by the hyperbolicity condition while the dynamics on the manifold is neutral (points are mapped onto themselves).

**Figure 3:**
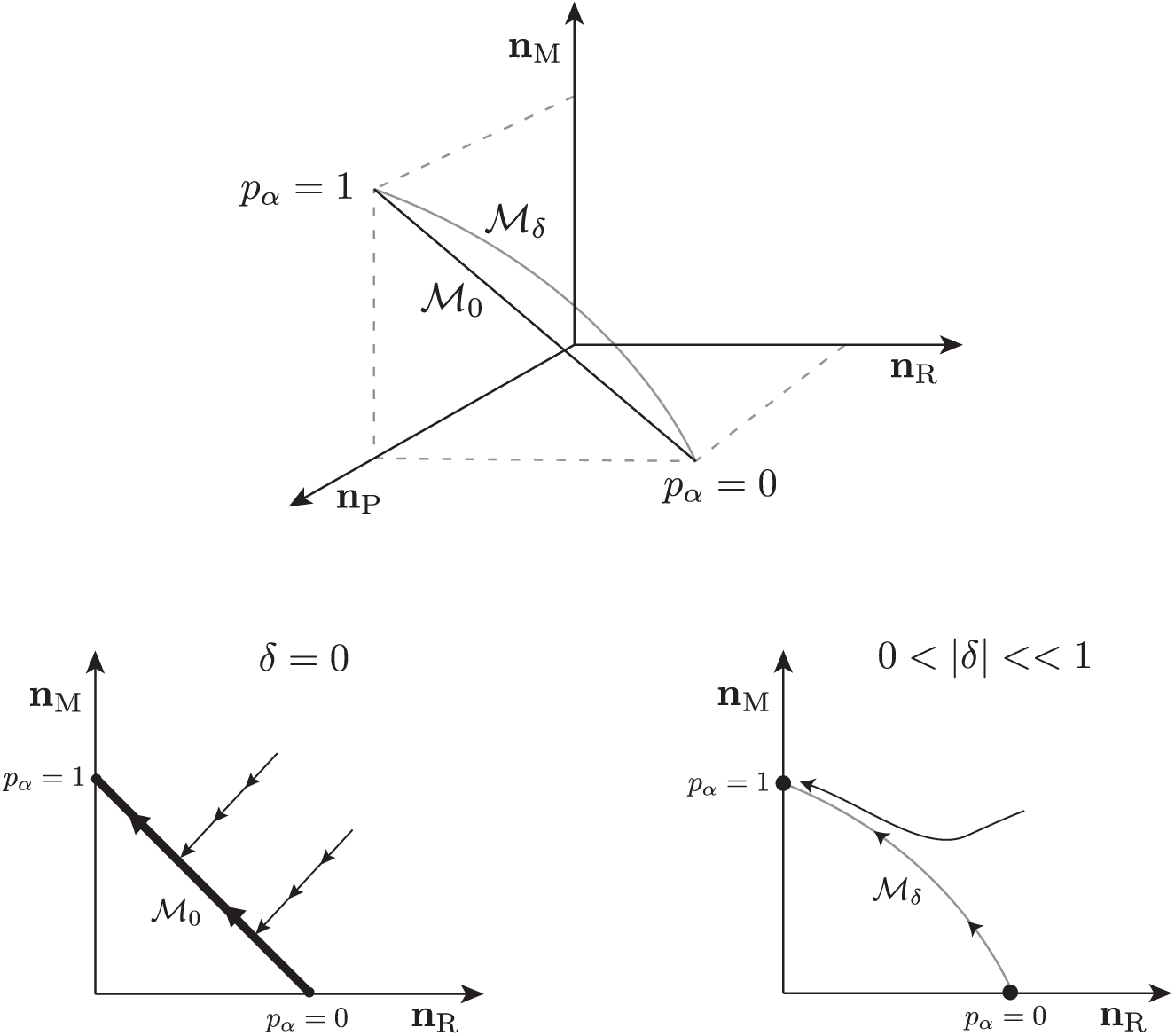
A diagram of the critical ℳ_0_ and slow ℳ_*δ*_ manifolds in the population state (phase) space where the axis are multidimensional and represent the phase space of mutants, residents and the rest of the community (if applicable) as given in the main text. **Top panel:** The critical manifold ℳ_0_ is obtained from the mutant-resident dynamics under phenotypic equality (*δ* = 0) by solving (21) and it consists of a line of equilibria. For (5) where *δ* is small but nonzero there exists a slow manifold ℳ_*δ*_, which is close to ℳ_0_ and has the same dynamical properties as ℳ_0_ (see bottom panels). **Bottom left panel:** The fast population (29) and slow evolutionary (30) dynamics of the singular system where *δ* = 0. The thin lines with arrows represent the fast dynamical convergence given by (29) to ℳ_0_ (where class-specific mutant frequencies are the weighted frequencies *p*_*α*_), and the thick line with arrows represents the slow evolutionary dynamics of *p*_*α*_ given by (30) on ℳ_0_ (in this example mutant frequency increases from 0 to 1). **Bottom right panel:** The mutant-resident dynamics (27) or (28) where *δ* is small but nonzero. The results of Fenichel (1979) say that since the (fast) dynamics for *δ* = 0 (bottom left panel) approaches ℳ_0_ so does the dynamics for small but non-zero *δ* approach ℳ_*δ*_. Moreover, the dynamics of *p*_*α*_ on ℳ_*δ*_ and its neighborhood, is equivalent of the (slow) dynamics of *p*_*α*_ on ℳ_0_ (left panel).

#### 3.2.2 Slow manifold

As elucidated above, the critical manifold ℳ_0_ is compact and normally hyperbolic, and therefore the results of Fenichel (1971, 1974, 1977, 1979, see also Appendix 6.4 and the references within) guarantee that a perturbed manifold ℳ_*δ*_, the so-called slow manifold (Hek, 2010; Jones, 1995), for the mutant-resident dynamics under phenotypic closeness exists, is close to, and has identical stability properties as ℳ_0_ (see also Figure 3, bottom panels). This slow manifold ℳ_*δ*_ is thus a set of points that are invariant under the dynamics of the full mutant-resident dynamics for small but nonzero *δ* (unlike in ℳ_0_, however, the points in ℳ_*δ*_ are not equilibria and hence the dynamical system maps them onto other points on ℳ_*δ*_), while in the neighborhood of ℳ_*δ*_ and ℳ_0_ the dynamics of the system (20) are equivalent. In other words, because the dynamics under phenotypic equality given by (20) approaches the critical manifold ℳ_0_, so does the dynamics under phenotypic closeness approach the slow manifold ℳ_*δ*_ (see Figure 3 and also, e.g., Jones 1995, Theorem 3, p. 62 and Theorem 6, p. 74). Moreover, the dynamics of *p*_*α*_ when restricted to ℳ_0_ (in slow evolutionary time) and the dynamics of *p*_*α*_ when restricted to ℳ_*δ*_ or to its neighborhood (in fast and slow time) are also equivalent (see a more detailed discussion in Appendix 6.4). This result plays a fundamental role in Section 4, where we prove the “invasion implies substitution”-principle by studying the singularly perturbed slow evolutionary dynamics of *p*_*α*_.

## 4 Invasion implies substitution in demographically class-structured ecological communities

We now prove the “invasion implies substitution”-principle for the population model of this paper whose resident dynamics is given in (3). We prove the principle by separating the timescales at which the various dynamical variables of the mutant-resident model (4) operate by using the weighted average mutant frequency *p*_*α*_. Because the dynamics of *p*_*α*_ is a function of class reproductive values ***α***, mutant frequencies **p**, total population densities **n** and the densities **n**_P_ of the other resident phenotypes and the rest of the ecological community, the complete mutant-resident dynamics for arbitrary phenotypic values 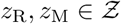 (*δ* arbitrary) can be written by extending (5) as:

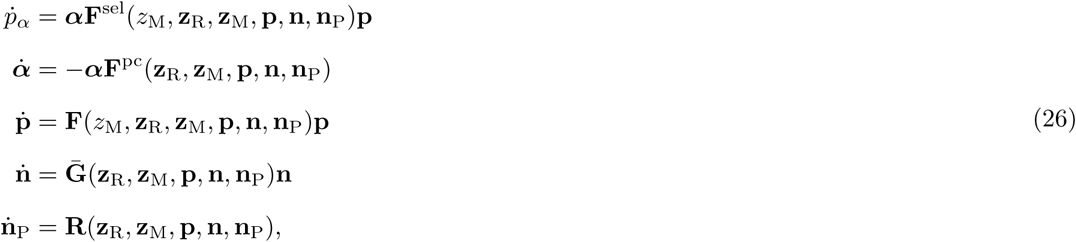

where we have for clarity included all the arguments. Next, we write the dynamics of (26) under phenotypic similarity in both fast and slow time, and then obtain two distinct limiting singular equations (by letting *δ* go to 0) that can be easily analyzed. Finally, we glue them back together by perturbing the obtained singular equations. By doing this the singular system (*δ* = 0) serves as an approximation to a mutant-resident dynamics under phenotypic similarity (*δ* small but nonzero) such that all its dynamical properties are preserved.

**Proof:** Let *t* denote the fast population dynamical time (the original time used throughout this paper) and let *τ* denote the slow evolutionary time (see also Figure 1). Setting *τ* = *δt* we obtain the relation *dτ* = *δdt* and then write the mutant-resident dynamics for closely similar phenotypes (*δ* small but nonzero) either using the original time variable *t*

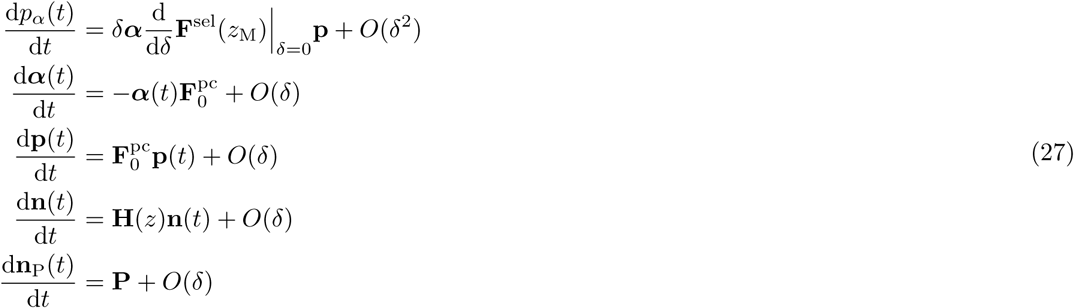

or using the new time variable *τ*

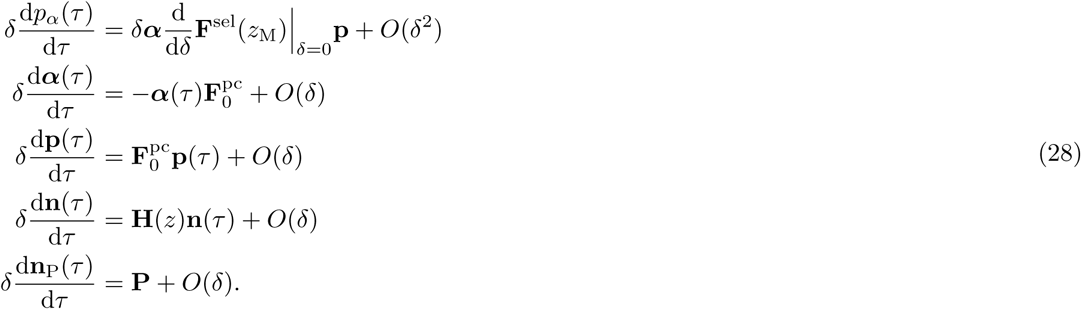

Since we haven’t yet taken any limits the two systems (27) and (28) are identical, the only difference is the notation. Let’s now take the limit *δ* → 0 and obtain two limiting singular equations, one for fast population dynamical time

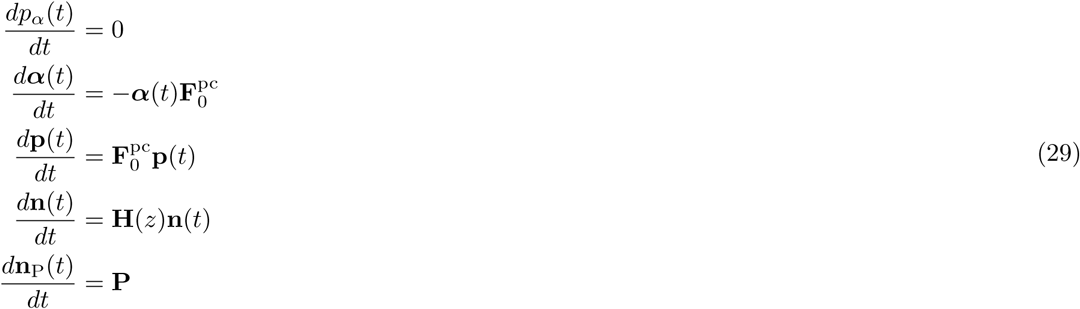

and the second for slow evolutionary time

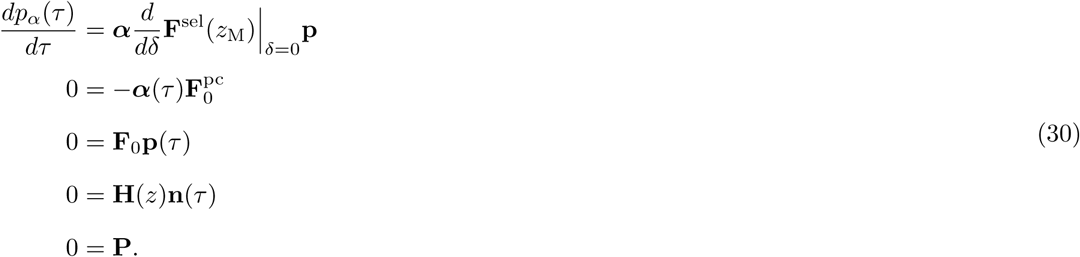

This confirms that in the fast population dynamical time (29) the average mutant frequency *p*_*α*_ stays constant and that the mutant-resident dynamics reaches the critical manifold ℳ_0_ as found in (25), and that the algebraic expression for ℳ_0_ can be obtained directly from (30).

Because the variables ***α*, p, n, n**_P_ in (30) have already reached the critical manifold ℳ_0_ and can thus be considered constant, we evaluate the right hand side of the first line in (30) at the ℳ_0_ to obtain

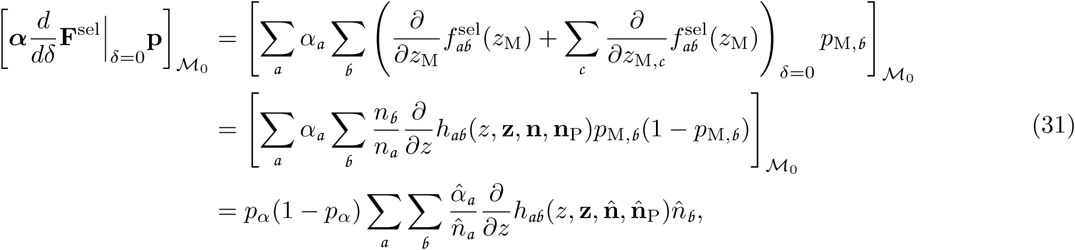

where we used (13). By using ***ν*** = (*ν*_*𝒶*_)_*𝒶*∈𝒟_ as a vector of reproductive values 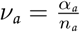 of an *individual* in class *𝒶* ∈ 𝒟 (see Section 3.1 and Appendix 6.5 for more details), then at ℳ_0_ we have

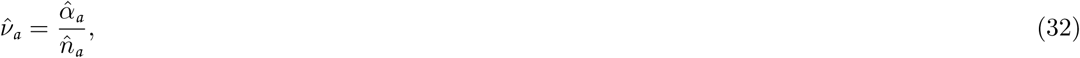

and using (31) we can write the slow (singular) mutant-resident evolutionary dynamics (30) with a single equation as

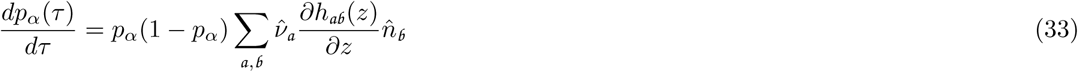

or in a matrix notation as

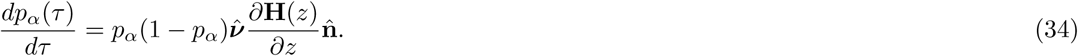

Alternatively, one can express (34) in terms of a probability distribution over all classes, i.e. in terms of class frequencies defined as 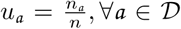, ∀*𝒶* ∈ 𝒟 where *n* = ∑_*𝒶*_ *n*_*𝒶*_ is the total population size. Because **u***n* = **n**, where **u** = (*u*_*𝒶*_)_*𝒶*∈𝒟_, one could also scale the individual reproductive values as **v** = ***ν****n* (Section 3.1 and Appendix 6.5) to get

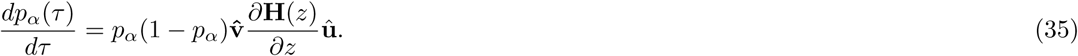

The two formulations (34) and (35) are equivalent, each providing a different perspective on the same evolutionary process. Because the matrix **H** gives the *individual* growth-rates, the expression in (34) describes how all (mutant) individuals in different classes contribute to the mutant evolutionary dynamics. In (35), the focus is on an average carrier of the mutant allele and how that representative individual contributes to the mutant dynamics when weighted over all classes the carrier of the mutant can be in.

Now, whichever formulation (34) or (35) is more convenient, geometric singular perturbation theory guarantees that after initial convergence, the mutant-resident dynamics (27)-(28) in the neighborhood of the manifold ℳ_*δ*_ is equivalent to (can be approximated by) the dynamics given by the two singular systems (29) and (30) (see Figure 3 and Appendix 6.4). In particular, the dynamics of the weighted mutant frequency *p*_*α*_ for small but nonzero *δ* near ℳ_*δ*_ can be approximated by the dynamics given in (34) and (35) (in Appendix 6.4 the Corollary 2 and the Section 6.4.4). We have thus proved the below “invasion implies substitution”-proposition and its Corollary, given the following assumption holds.

### Assumption (A).

*Assume that the resident ecological community as defined in* (3) *contains a hyperbolically stable equilibrium* 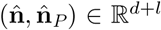*to which the resident population converges to and then stays at.*

**Invasion implies substitution-principle.** *Consider an ecological community with a polymorphic demographically structured population as defined in* (3), *and assume that (A) holds. Suppose that one of the alleles in the population undergoes a mutation, and that the resulting mutant phenotype* 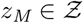 *and its ancestral resident phenotype* 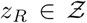 *are closely similar, i.e. δ* = *z*_*M*_ − *z*_*R*_ *for some small δ* ≠ 0. *Then, for sufficiently large time t and/or small δ, the dynamics of the weighted mutant frequency p*_*α*_ *in the resulting mutant-resident ecological community* (26) *can be approximated on the original time scale by*

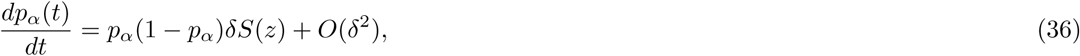

*where the frequency-independent selection gradient S*(*z*) *can be expressed as*

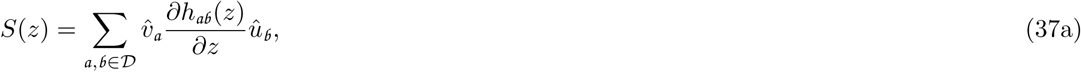

*or in matrix notation as*

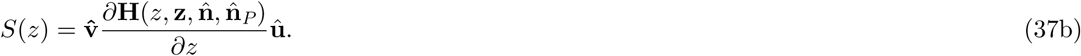

*Successful invasion of a mutant implies the substitution of the resident.*

### Corollary (C).

*The subset of* 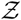 *where the assumption (A) holds and where the selection gradient* (37) *is nonzero, S*(*z*) ≠ 0, *indicates the set of phenotypes for which the “invasion implies substitution”-principle holds and where the phenotype is under directional selection.*

## 5 Discussion

We proved positive answer to all three questions (I)-(III) delineating the “invasion implies substitution”-principle program (see Introduction) for scalar-valued, polymorphic and well-mixed haploid reproducing populations that are part of a larger ecological community and that are structured into finitely many demographic classes.

### 5.1 The separation of population dynamical and evolutionary variables

We proved the “invasion implies substitution”-principle by separating the population dynamical and evolutionary timescales using the weighted average mutant frequency, and then singularly perturbed the mutant-resident dynamics given as ordinary differential equations (Fenichel, 1979; Wiggins, 1994; Jones, 1995; Hek, 2010; Kuehn, 2015; Dercole and Geritz, 2016) using the phenotypic deviation *δ* as the perturbation parameter. In this method, which is fully detailed in Appendix 6.4 for the present context, one proceeds in three steps. First, one must be able to write the mutant-resident dynamics for small values of *δ* in a fast-slow form 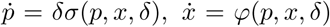, where *p* represents a weighted mutant frequency in the population and *x* should capture all the fast (population dynamical) variables. In Section 3, however, it became apparent that for small *δ* all dynamical variables are fast variables, including class-specific and mean mutant frequencies, and so the model could not readily be written in the above fast-slow form. The solution here was to introduce a new variable which operates purely in slow evolutionary time and is a proxy for the mutant frequency. In Section 3.1 we showed that such a variable is the average mutant frequency weighted by class reproductive values (Taylor, 1990; Leturque and Rousset, 2002; Rousset, 2004; Lehmann and Rousset, 2014; Engen et al., 2014; Lehmann et al., 2016).

Once the mutant-resident dynamics is in the fast-slow form, in the second step one starts analyzing the dynamics of the weighted mutant frequency *p*. Because studying its dynamics for nonzero *δ* is a complicated task, one hopes that the dynamics of the much easier model where *δ* = 0 (i.e., the neutral model) could serve as an approximation for small but nonzero *δ*. To achieve this, one must first scale time by using *δ* as the scaling parameter and then write the mutant-resident dynamics in both fast *t* and slow time *τ* = *δt* while letting *δ* go to zero. In this step one thus analyzes two singular systems, one in fast time where *p* is constant and *x* fluctuates according to 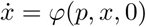, and the other in slow time where *x* is constant (i.e. is at the steady state) and *p* fluctuates according to 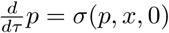. For us to be able to draw conclusions from this singular system the variable *x* must converge to its steady state in fast time. In our model this follows directly from the assumption that the resident steady state 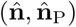 is hyperbolically stable, i.e. the real part of all eigenvalues of the Jacobian of the linearized resident dynamics are all negative.

In the third and final step one perturbs the above singular equations by applying geometric singular perturbation results for ordinary differential equations developed in Fenichel (1979). Provided certain conditions are satisfied, one can then equate the dynamics of the singular equations where *δ* = 0 with the original system where *δ* is small but nonzero (i.e. the perturbed system). Conveniently, the sufficient condition for such a singular perturbation to be possible is that the steady state is hyperbolic, which is true by assumption. Therefore, if invasion implies substitution holds for the singular system, it holds also for the original (perturbed) mutant-resident dynamics whenever the steady state is hyperbolic.

The above-mentioned procedure can be applied to more generally structured population models than the one presented in this paper. First of all, the singular perturbation results in Fenichel (1971, 1974, 1977, 1979) allow a direct generalization of our result to models with attractors other than equilibria, e.g. to limit cycles where population experiences deterministic periodic fluctuations. Because including more complicated attractors would require some amount of additional notions (e.g. time-dependent reproductive values as e.g. discussed in Lion, 2018b) we choose to leave this generalization for future work. Second, more recent but similar results on invariant manifolds for semiflows (Bates et al., 1998, 2000; Pliss and Sell, 2001) accommodate infinite-dimensional population structure e.g. continuous age or size distributions. However, calculations of the hyperbolicity of steady states is considerably more involved in such cases (Greiner et al., 1994; Kuehn, 2015; Cantrell et al., 2017) and directly applicable only to models where the transitions between spatial or demographic classes is density-independent (Greiner et al., 1994; Cantrell et al., 2017).

### 5.2 Selection gradient as a map between ecology and evolution

The expression for the selection gradient (37) was obtained directly from the timescale separation argument given in Section 4. We found that the selection gradient can indeed be written as conjectured in (2), but without class-specific trait expression, since we considered only a single scalar trait for simplicity of presentation (and the extension to class-specific trait is direct), and with relatedness **r** yet playing no role. Relatedness plays no role because we assume infinitely large population sizes with no spatial structure (i.e., a well-mixed population) and hence genealogical relationships between any two individuals do not affect the direction of selection. Nevertheless, the selection gradient can be written solely in terms of resident population dynamical variables and resident growth-rates. This is practical since one can then calculate directly from the resident dynamics which mutations can and cannot fix into the population, that is, one can calculate the fate of the mutation before the mutation actually takes place. In this sense, the selection gradient is a “map” from the ecological to the evolutionary model (see Figure 1). Moreover, it is a map that collapses the potentially multi-dimensional population structure into a scalar-valued measure.

An analogous selection gradient for a model that has the same biological scope as in the present paper has been previously derived in Lion (2018a,b). The model and the method obtaining the selection gradient however depart from ours in that in Lion (2018a,b) the polymorphism is assumed tightly clustered around its mean and that the dynamical equations were formulated in terms of change in mean phenotype. Such a formulation provides links between the dynamics of the mean trait value and the “invasion implies substitution”-principle and is thus complementary to our approach. The drawback in this approach, however, is that the timescale of dynamical variables such as class-specific mutant frequencies is not easily accessible. Consequently, in particular our results on the critical manifold ℳ_0_ (Section 3.2), allows us to confirm that as the class-specific trait variance is proportional to the class-specific mutant frequencies, it is indeed a fast variable approaching the population mean trait variance, a result that was left open in Lion (2018a). We conjecture that the ideas on tightly clustered phenotypes developed in Meszéna et al. (2005) together with the results derived in this paper fully justify the selection gradient presented in Lion (2018a,b).

### 5.3 Selection gradient as a map between fitness landscape and the direction of evolution

The main implication of the “invasion implies substitution”-principle is that it indicates the set of resident phenotypes that can be invaded by a mutant phenotype (or more precisely, invaded by a phenotypic deviation *δ*), thus providing a tool to study the *meso*-evolutionary dynamics of the trait under selection (panel C in Figure 1 and Corollary in Section 4). The sequential invasion and substitution can occur whenever the steady state is hyperbolic, thus excluding the possibility of bifurcations that may lead to catastrophic extinctions, and whenever the selection gradient *S*(*z*) is nonzero, i.e. as long as we are away from the extrema of the adaptive landscape (indicated with grey circles in panel C Figure 1). Such extrema identify the phenotypic values where invasion no longer implies substitution and where more complicated evolutionary behaviour can occur (Geritz et al., 1998; Priklopil, 2012; Dercole and Geritz, 2016). Nevertheless, because we have formulated our model for arbitrarily polymorphic resident populations, the “invasion implies substitution”-principle holds *whenever* the selection gradient is non-zero (and the steady state is hyperbolic). This is even true after the evolutionary dynamic converges and escapes a phenotypic value that is a branching point: “invasion implies substitution”-principle still governs the direction of evolution after the appearance of new morphs and thus over the whole fitness landscape (except near points with measure zero). Yet, in the presence of evolutionary branching, a selection gradient is needed for each branch as depicted in Figure 1.

### 5.4 Conclusion

This study is part of a quest aiming at generalizing and formalizing the hypothesis that traits under frequency and/or density dependent selection are generically subject to directional gradual change, whenever mutations cause only small deviations to the phenotype under selection (and in the absence of genetic constraints). Further, directional selection should be quantifiable by a selection gradient that consist of reproductive value and relatedness weighted fitness differentials. We here confirmed this hypothesis for well-mixed ecological communities with demographically (physiologically) class-structured populations. Our results are directly applicable to several well-known models, such as SIR-models in epidemiology and stage-structured models in life-history studies, and will be generalized to spatially structured population with limited dispersal in a forthcoming study.

## 5.5 Acknowledgments

TP was supported by the Swiss NSF grant PP00P3-123344 to LL.

## 6 Appendix

### 6.1 Relative growth-rate for arbitrary phenotypes

In the main text, we derived the dynamics for class-specific mutant frequencies (5)-(8), where we obtained a partition for the relative growth-rate matrix for a (single) mutant **F**(*z*_M_) = **F**^sel^(*z*_M_)+ **F**^pc^, with a term **F**^pc^ that is independent of the phenotype of the (single) mutant. Here, we confirm that such a partition exists independently of the phenotype of the individual whose relative growth-rate we are considering by proceeding the same way as in the main text, except that we do not specify the phenotype of the individual whose relative growth-rate we are calculating. That is, we have

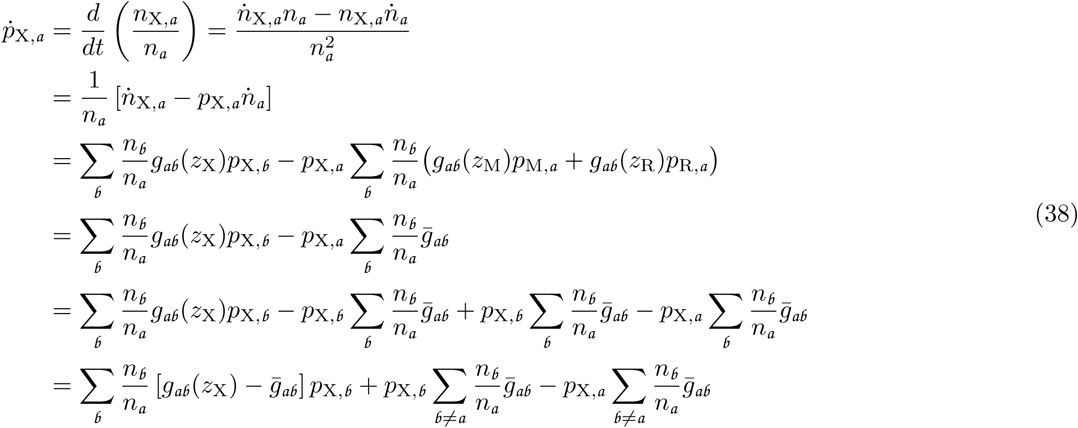

for all *𝒶* ∈ 𝒟, where 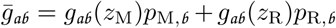 and *X* ∈ {M, R}. Defining **p** := **p**_M_ and **1** − **p** := **p**_R_ as the vector of class-specific mutant and resident frequencies, respectively, we can write

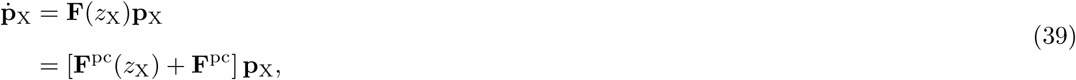

where the entries of **F**^pc^(*z*_X_) and **F**^pc^(*z*_X_), respectively, are

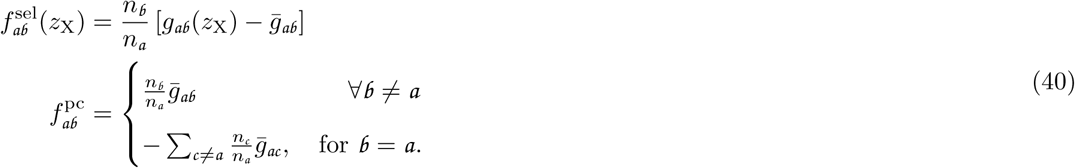

Notice that the component that gives the rates at which passive changes occur **F**^pc^ is the same for both mutant and resident phenotypes. In fact, an analogous expression can be derived for any polymorphism as long as 1 = ∑_X_ *p*_X,*𝒶*_ for all *𝒶* ∈ 𝒟.

### 6.2 Mean mutant frequency *p*_M_ and the dynamics of class frequencies

In the main text, we showed that class-specific mutant frequencies **p** are both fast and slow dynamical variables. More precisely, we showed that under phenotypic equality (*δ* = 0) the dynamics is dominated by the terms of order *O*(1) (Section 3) and that **p** approaches a line of equilibria where 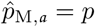 for all *𝒶* ∈ 𝒟, after which the dynamics is dominated by the terms of order *O*(*δ*) along this line of equilibria (Section 3.2). Here, we confirm that the same applies for the mean (arithmetic) mutant frequency in the total population *p*_M_ = **up** = ∑_*𝒶*_ *u*_*𝒶*_ *p*_M,*𝒶*_.

To confirm this, it is sufficient to show that **u** approaches in fast population dynamical time an isolated equilibrium which persist under perturbation of *δ*. If this is so, then the dynamics of *p*_M_ is first dominated by the terms of order *O*(1) and then of order *O*(*δ*) and we get our claim. This is checked immediately from the following Section 6.2.1 where we detail the dynamics of class frequencies **u**: because by assumption the steady state 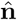 is hyperbolic so is the steady state 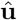 in (49) (and thus it persists under perturbations).

#### 6.2.1 The dynamics in terms of class frequencies

In this section we will re-write the resident dynamics (3) and the relative mutant-resident dynamics (5) in terms of total population densities and class frequencies, which are respectively defined as

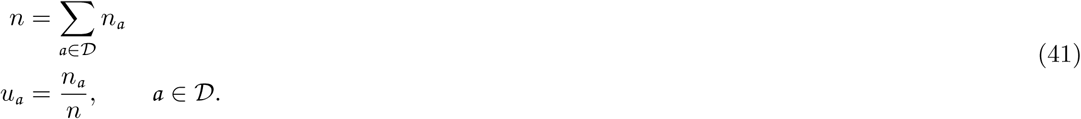

Note that since *n* is a scalar we have the relation

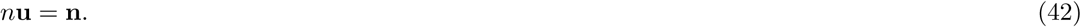

##### Resident dynamics

The dynamics of the total density is obtained by using (41) and by differentiation

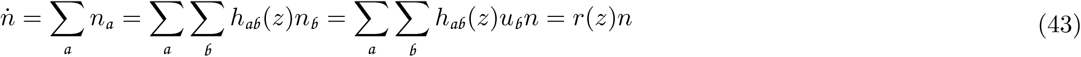

where *r*(*z*) := *r*(*z*, **z**, *n*, **u, n**_P_) = ∑_*𝒶*_ ∑_*𝒷*_ *h*_*𝒶𝒷*_ (*z*)*u*_*𝒷*_ is the total mean growth-rate of an individual in the resident population. The dynamics of class frequencies is obtained by using (41), (43), and by differentiation

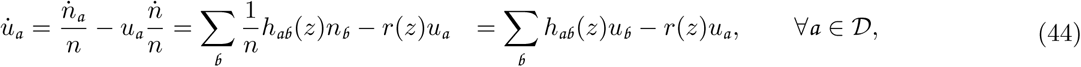

or in a matrix notation

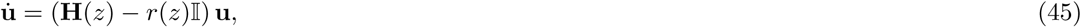

where 𝕀 is the identity matrix. We have thus obtained that the resident dynamics (3) can be rewritten as

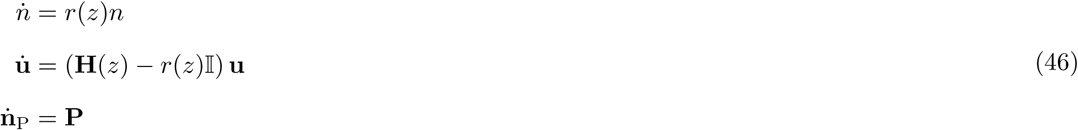

##### Relative mutant-resident dynamics

Using (5) and an analogous derivation to the previous section, we obtain for the relative mutant-resident dynamics

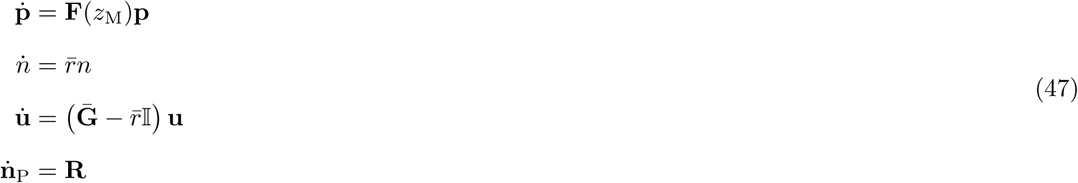

where 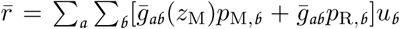 is the total mean growth-rate of an individual in the total population. Notice that alternatively 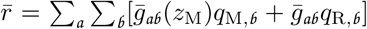, where 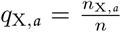 is the probability that given an individual is sampled from the total population it is an individual in class *𝒶* ∈ 𝒟 with phenotype *z*_X_ ∈ {*z*_M_, *z*_R_}.

##### Steady state under phenotypic equality

In section 3.2, we found that the steady state 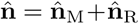 under phenotypic equality *δ* = 0 can be solved from

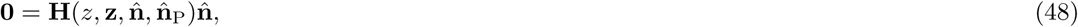

and is thus the right eigenvector of the resident matrix 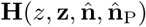 associated with the eigenvalue 0. Here, we are interested to express the steady state under phenotypic equality in terms of the total population size *n* and class frequencies **u**. Using Section 2.4, we have

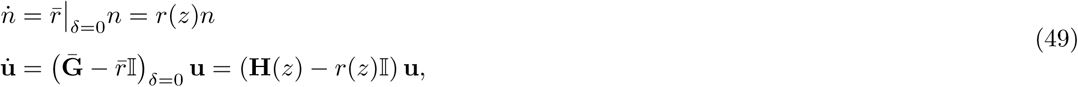

where the (non-trivial) solutions 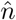 and 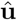 are obtained from

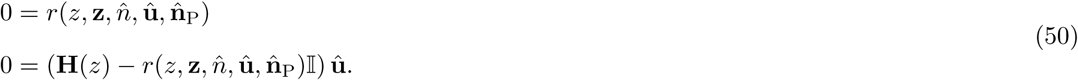

Using (42) the steady state can be written as

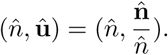

Note that because *r*(*z*) in-front of the identity matrix in (49) is scalar-valued, both 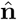 and 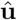 are the right eigenvectors of 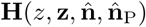 associated with the eigenvalue 0 (this is in fact obvious since we have the relation *n***u** = **n**, i.e. an eigenvector scaled by a scalar is also an eigenvector associated with the same eigenvalue).

### 6.3 Infinite number of equilibria in ℳ_0_

Here, we give an argument as to why the singular system *δ* = 0 contains infinite number of equilibria. Because 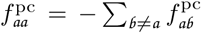 the matrix **F**^pc^ is a transition matrix with an eigenvalue 0. Because the eigenvalue is solved from 0 = det[**F**^pc^ − 0 · 𝕀] = det[**F**^pc^], the determinant of **F**^pc^ is zero implying that it is not an invertible matrix and hence **F**^pc^**p** doesn’t have a unique isolated solution **p** (see e.g. Hirsch et al., 1974, Proposition on p. 80).

### 6.4 Fenichel’s Theorems

Here, we go in detail through the results of Fenichel (1971, 1974, 1977, 1979) that are relevant for the “invasion implies substitution”-principle so that our paper is self-contained. This section can be seen as a general recipe on how to translate any mutant-resident dynamical system (that is expressed in terms of ordinary differential equations) into a singular perturbation problem, and how the theory of Fenichel allows us to obtain a complete description of the dynamics for the mutant frequency in the full mutant-resident model where *δ* is small but nonzero. We will in most part follow the exposition of Jones (1995) and Hek (2010) (and with a small dose of Kuehn, 2015).

The full mutant-resident dynamical model (arbitrary *δ*) as given in (26) is our starting point

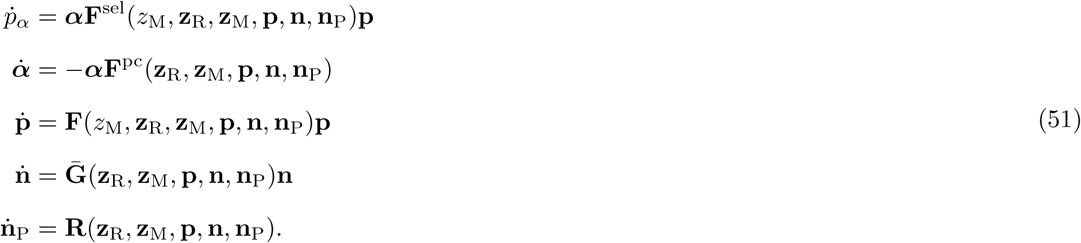

This system can be equivalently written as

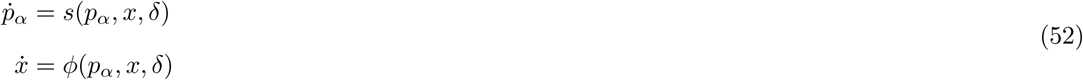

where 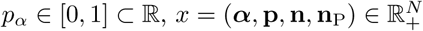 (with *N* = 3*d* + *l*),and where

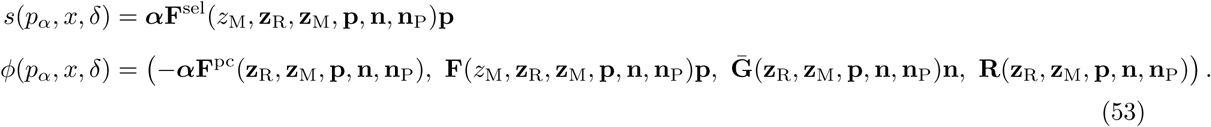

***(H1)*** The functions *s, ϕ* are sufficiently smooth.

Given (H1) the Taylor expansion of (52) about *δ* = 0 is

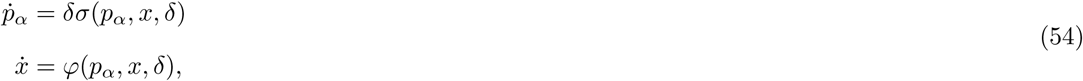

where

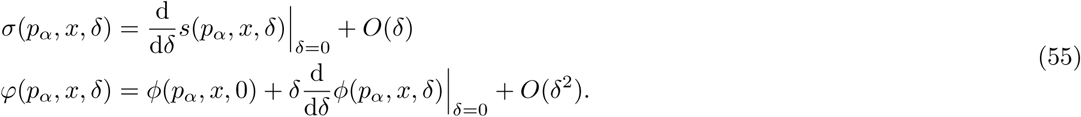

#### 6.4.1 The relative mutant-resident dynamics

As in the main text, let *t* be the fast (population dynamical) time and *τ* = *δt* the slow (micro-evolutionary) time. For simplicity we will use a dot for the time derivative in fast time (as in the main text) and a comma for the time-derivative in slow time.

##### The original (perturbed) fast and slow system

We can write the system (53), for small but nonzero *δ* using (54), in both fast and slow time, respectively, as

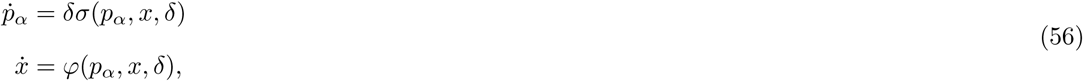

and

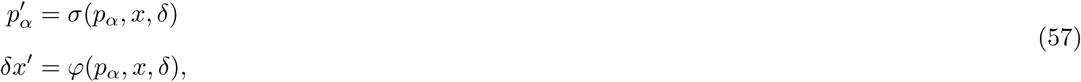

and we re-iterate that *p*_*α*_ ∈ [0, 1] ⊂ ℝ and 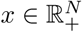 (with *N* = 3*d* + *l*).

##### The singular fast and slow system

By taking the limit *δ* → 0 and by applying (55) we obtain two singular sets of equations for both fast and slow time:

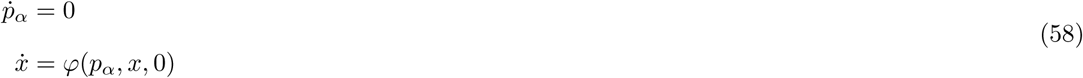

and

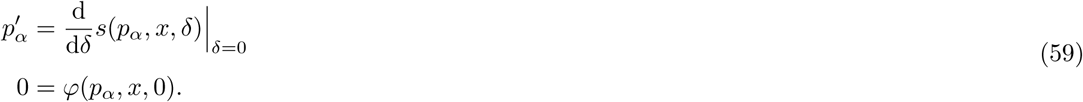

#### 6.4.2 Fenichel’s Theorems 1 and 2

Throughout, whenever we are referring to a distance between two nonempty sets we use the notion of Hausdorff distance (see e.g. Kuehn, 2015, p. 55).

##### Critical manifold

The set of critical (equilibrium) points *φ*(*p*_*α*_, *x*, 0) = 0 is obtained by solving *N* equations yielding an 1-dimensional manifold. That is, the set of critical (equilibrium) points is parametrized by *p*_*α*_. We will denote the biologically relevant subset of those points with

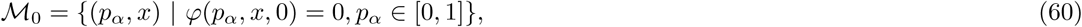

which is the critical manifold mentioned in the main text (recall eq. 25).

***(H2)*** The set ℳ_0_ is compact and normally hyperbolic. Moreover, the linearization of (56) at each point in ℳ_0_ has exactly one eigenvalue on the imaginary axis and *N* eigenvalues on the left-side of the imaginary axis (i.e. the manifold ℳ_0_ is locally asymptotically stable; Jones, 1995, p. 49).

The normal hyperbolicity of the manifold ℳ_0_ means that the dynamics (or flow; henceforth we will use both words depending which one is more descriptive) in the neighborhood of this manifold is governed by the non-zero eigenvalues and the flow on the manifold is governed by the zero eigenvalue, i.e. the flow on the manifold is neutral (each point is mapped to itself, i.e. each point is invariant under the flow).

The following theorem is an adaptation from Jones (1995, Theorem 1, p. 49) and Hek (2010, Theorem 2, p. 354).

###### Fenichel’s Invariant Manifold Theorem 1.

*Assuming (H1) and (H2), for δ non-zero but sufficiently small, there exists a (slow) manifold* ℳ_*δ*_ *that lies within O*(*δ*) *of* ℳ_0_ *and is diffeomorphic (“isomorphic”) to* ℳ_0_. *Moreover, it is invariant under the flow of* (56).

This theorem implies that the restriction of the flow of (56) to ℳ_*δ*_ is a small perturbation of the flow of the limiting (or singular) problem (59). This can be directly seen in the case where ℳ_0_ is given by a graph of a function *π*_0_(*p*_*α*_) (which can be done at least locally because ℳ_0_ is normally hyperbolic and hence one can apply the Implicit Function Theorem), where the subscript 0 indicates that we are discussing the limiting (singular) problem where *δ* = 0. If

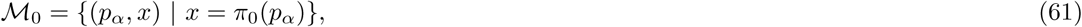

then the slow manifold ℳ_*δ*_ can be represented as a small perturbation *π*_*δ*_ of *π*_0_ as

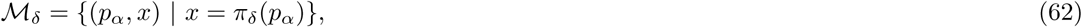

where the subscript *δ* in *π*_*δ*_ indicates that we are discussing the original perturbed problem (56) - (57) for non-zero but small *δ*. Substituting (62) into the equation for slow time (57) we obtain

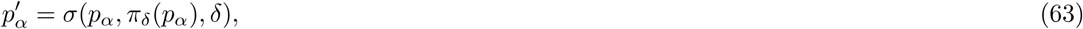

which indeed reduces to the limiting (singular) problem 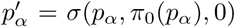 as in (59) by taking the limit *δ* → 0 (see also Hek, 2010, the final paragraph on p. 354).

At the moment we only know about the flow on (or restricted to) the manifold ℳ_*δ*_ but not in the neighborhood of ℳ_*δ*_. This will be addressed in the following theorem, but before presenting it we must introduce some additional notation in order to define the notion of a stable manifold of a set. First, recall that the critical manifold ℳ_0_ in our model consists, by definition, of a set of hyperbolically stable critical points {*y* = (*p*_*α*_, *x*)} (there are no unstable points, see (H2)). Then, the so-called stable manifold theorem guarantees that any such equilibrium point *y* ∈ ℳ_0_ has a stable manifold *W* ^s^(*y*) associated to it (see e.g. Wiggins, 1994). The stable manifold of *y* is the set of points from which the dynamics converges to *y* under the flow of (56) (that is, roughly speaking, *W* ^s^(*y*) is the basin of attraction of the point *y*). Because *x* is *N* -dimensional in our model, each manifold *W* ^s^(*y*) is an *N* -dimensional manifold in the phase space 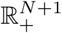. Then,

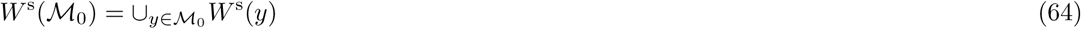

is the *N* + 1-dimensional stable manifold of the set ℳ_0_ (see also Hek, 2010, p. 372).

With this notation, we can state the following theorem, which is an adaptation from (Jones, 1995, Theorem 3, p. 62) and (Hek, 2010, Theorem 4, p. 359).

###### Fenichel’s Invariant Manifold Theorem 2.

*Assuming (H1) and (H2), for δ non-zero but sufficiently small, there exists a manifold W*^*s*^(ℳ_*δ*_) *that is diffeomorphic (“isomorphic”) to and lies within O*(*δ*) *of W*^*s*^(ℳ_0_). *Moreover, W*^*s*^(ℳ_*δ*_) *is invariant under the flow of* (56).

Theorem 2 proves the existence of a perturbed smooth manifold *W* ^s^(ℳ_*δ*_) whose points converge towards the invariant manifold ℳ_*δ*_ at an exponential rate forward in time under the flow (56). That is, the manifold *W* ^s^(ℳ_*δ*_) is stable as the name suggests. However, and this is important to note, *W* ^s^(ℳ_*δ*_) is stable in a *different sense* than *W* ^s^(ℳ_0_). This is because whereas every point *y*_0_ on ℳ_0_ is invariant (each point maps to itself under the flow, i.e. they are equilibrium points) and *W* ^s^(*y*_0_) is invariant under the flow that approaches *y*_0_, points *y*_*δ*_ on ℳ_*δ*_ are *not* invariant (they are not equilibria). The question is thus in which sense the manifold *W* ^s^(ℳ_*δ*_) is stable. This information is essential, because in principle it is possible that the dynamics in *W* ^s^(ℳ_*δ*_) approaches a specific point on ℳ_*δ*_ (say *p*_*α*_ = 1), but the dynamics *on* ℳ_*δ*_ approaches a different point (say *p*_*α*_ = 0). This thus raises the question of the condition under which one can equate the dynamics of *p*_*α*_ on ℳ_0_ in the singular model to the dynamics of *p*_*α*_ *near* ℳ_*δ*_ in the original model with small but non-zero *δ*? This question is addressed in the next section.

#### 6.4.3 Fenichel’s Theorem 3 and its Corollaries

We now restrict our attention to a small neighborhood *D* of ℳ_*δ*_, where we can safely assume that the eigenvalues of the linearization of (56) dominate the dynamics, and focus on the trajectories in *W* ^s^(ℳ_*δ*_) that are in the neighborhood *D*. For this, let *y* · *t* denote the state of the dynamical system (i.e., a point 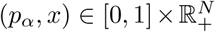) resulting from the application of the vector field (56) for a length of time *t* to initial state *y* (thus, *y* · *t* can be thought of as the solution of the system at time *t* given initial condition *y*), and let *y* · [*t*_1_, *t*_2_] denote the resulting trajectory over the interval [*t*_1_, *t*_2_] (Hek, 2010, Section 6.1.). Similarly, let *A* · *t* denote the set of states resulting from the application of the dynamical system eq. (56) for a length of time *t* to the set of states *A*.

The following definition is an adaptation from Hek (2010, Definition on p. 376) and Jones (1995, Definition 3 on p. 74).

##### Definition

The forward evolution of a set *A* ⊂ *D* restricted to *D* is given by the set

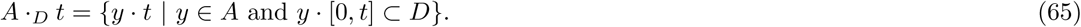

The following version of the theorem is an adaptation from Jones (1995, Theorem 6, p. 74) and Hek (2010, Theorem 8, p. 376 and Figures in Section 6.1.).

##### Fenichel’s Invariant Manifold Theorem 3.

*Assume (H1)-(H2). For every y*_*δ*_ ∈ ℳ_*δ*_, *there exists an N-manifold*

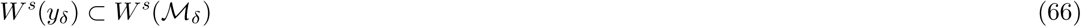

*that is O*(*δ*) *close and diffeomorphic to W*^*s*^(*y*_0_). *The family* {*W*^*s*^(*y*_*δ*_) | *y*_*δ*_ ∈ ℳ_*δ*_} *is invariant in the sense that*

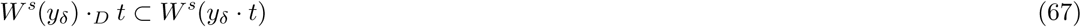

*if y*_*δ*_ · *r* ∈ *D for all r* ∈ [0, *t*].

The main point to notice here is that while a manifold *W* ^s^(*y*_*δ*_) is not invariant under the flow, and hence is called a *fibre* rather than a stable (invariant) manifold as e.g. is *W* ^s^(*y*_0_) (Jones, 1995, Section 3.3.), the family of such fibers is invariant under the flow. This somewhat abstract theorem is depicted in Figure 4 (adopted from two diagrams in Hek (2010, p. 377).

**Figure 4:**
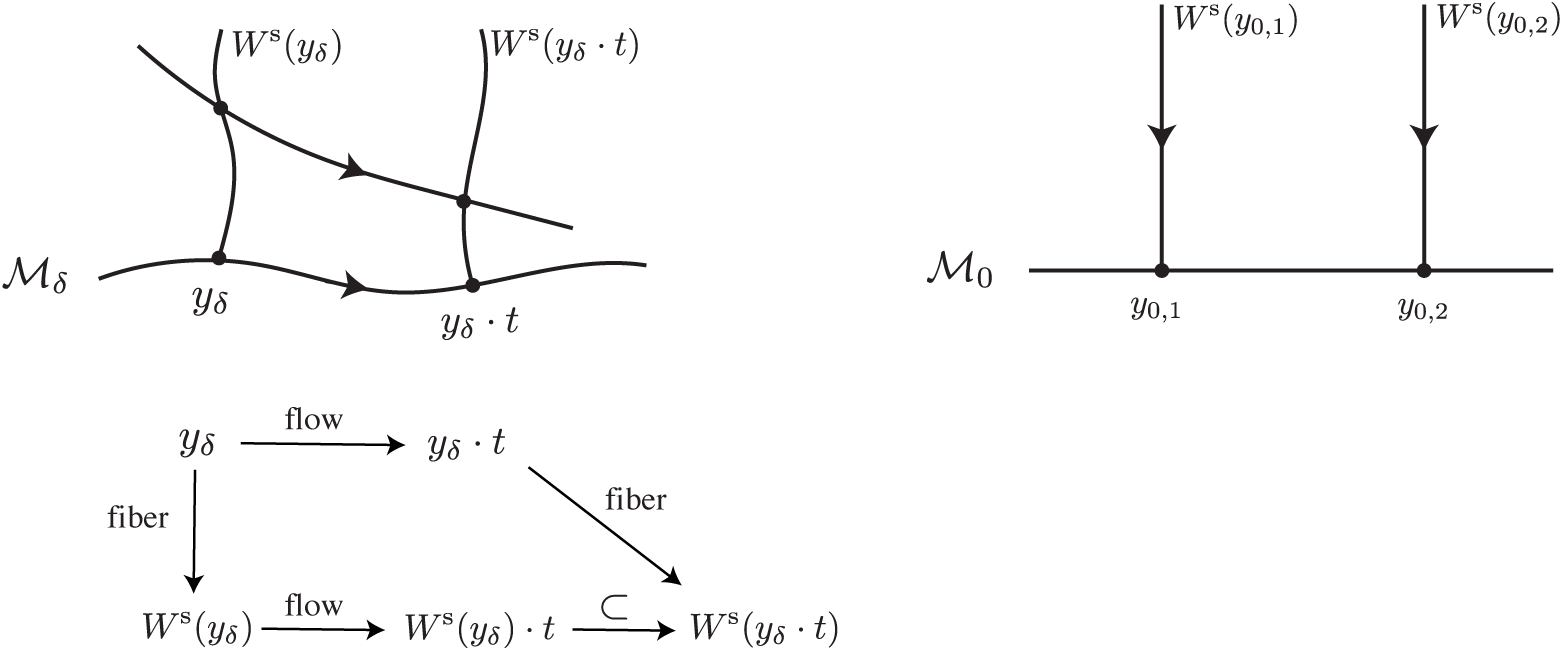
**Left diagrams** depict the Fenichel’s Theorem 3. Upper diagram shows the slow manifold ℳ*δ* and two fibers *W*^s^(*y*_*δ*_) and *W* ^s^(*y*_*δ*_ · *t*) that are attached to points *y*_*δ*_ ∈ ∑_*δ*_ and *y*_*δ*_ · *t* ∈ _*δ*_, and a trajectory that traverses them. As seen from the diagram, the fibers *W*^s^(*y*_*δ*_) and *W*^s^(*y*_*δ*_ · *t*) are not invariant under the flow. The bottom diagram presents an abstraction of the upper diagram on forward evolution of points on ℳ_*δ*_ and its neighborhood. **Right diagram** is for comparison and depicts the critical manifold ℳ_0_ and the stable manifolds *W*^s^(*y*_0,1_), *W*^s^(*y*_0,2_) attached to points *y*_0,1_ ∈ ℳ_0_ and *y*_0,2_ ∈ ℳ_0_, respectively.

The following version of the corollary is an adaptation from Jones (1995, Corollary 1, p. 76) and Hek (2010, Corollary 9, p. 377).

##### Corollary 1.

*There are constants κ, β* > 0 *so that if y* ∈ *W*^*s*^(*y*_*δ*_) ∩ *D, then*

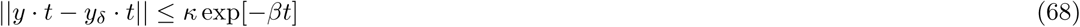

*for all t* ≥ 0 *for which y* · [0, *t*] ⊂ *D and y*_*δ*_ · [0, *t*] ⊂ ℳ_*δ*_.

The following corollary immediately follows (Hek, 2010, see the discussion after Corollary 9, p. 378):

##### Corollary 2.

*Suppose that y* ∈ *W*^*s*^(ℳ_*δ*_) *has a base point y*_*δ*_ ∈ ℳ_*δ*_, *then*

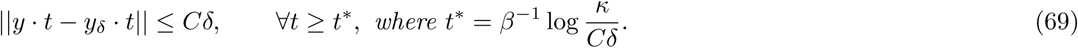

This means that we can find a point in time *t*^∗^ after which the distance between any two points, one point on ℳ_*δ*_ and the other in the neighborhood *D*, is of distance *O*(*δ*). Corollary 2 therefore gives the justification for the bottom right panel of Figure 3: a trajectory *y* · [*t*^∗^, *t*] in the neighborhood of ℳ_*δ*_ is approaching a trajectory *y*_*δ*_ · [*t*^∗^, *t*] on ℳ_*δ*_. Note the resemblance of the inequality in Corollary 2 to Nagylaki (1979, equation (37) on p. 440).

#### 6.4.4 Relating the slow-time singular system (34)-(35) to Equation (1)

From Theorem 1, we obtained that ℳ_*δ*_ and ℳ_0_ are *O*(*δ*)-distance away, that is, with some abuse of notation, we got

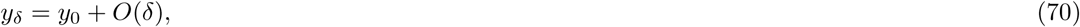

which in the model (51) reads as

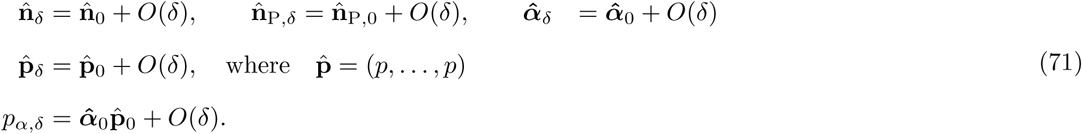

Here, the subscript *δ* and 0 denote that those variables take values on ℳ_*δ*_ and ℳ_0_, respectively. Then, using Corollary 2, we have that the estimates (70) and (71) hold 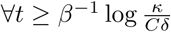, i.e.

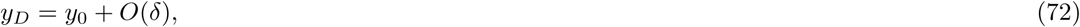

which in the model (51) reads as

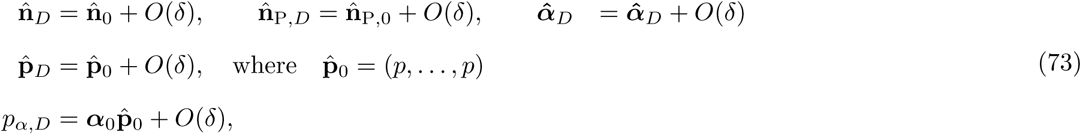

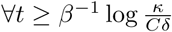, where we use the subscript *D* to denote that *y* takes a value in the small neighborhood *D* of ℳ_*δ*_. Moreover, by substituting (73) into *∂***H**(*z*)*/∂z* we also have 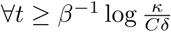 that

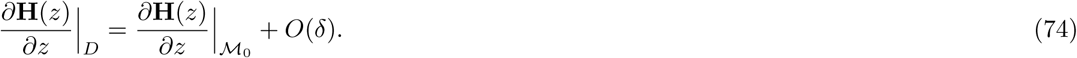

Now, in the main text (34) (or alternatively (35)) we derived for the slow singular system *δ* = 0 an equation

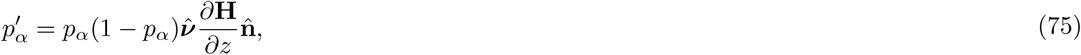

and by using (73)-(74) and the Corollary 2 we have that *p*_*α*_ in the neighborhood *D* of ℳ_*δ*_ can be written in slow evolutionary time *τ* as

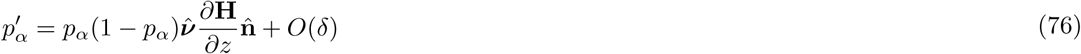

and therefore in fast original time *t* as

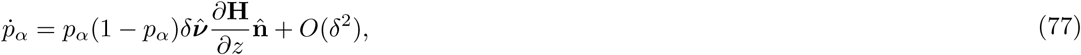

whenever 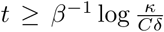 (which can also be written in terms of *τ*). This gives a full justification to equations (1) and (36) in the main text.

### 6.5 Individual reproductive values

This exposition in this Appendix is motivated by Lion (2018b) who, in contrast to standard practice in calculating the reproductive values only at the steady state, defined the reproductive values in both fast population dynamical as well as slow evolutionary time. We, however, depart from the exposition of Lion (2018b) by deriving a dynamical equation (analogues to the one in Lion (2018b)) for an alternatively scaled definition for individual reproductive value. Moreover, in the final Section 6.5.4 we show an alternative derivation for the dynamics of the weighted mutant frequency *p*_*α*_ by using such individual reproductive values.

In e.g., Taylor (1990); Lion (2018b) the individual reproductive values are defined as

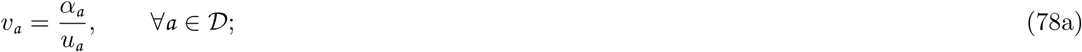

namely, such that they satisfy the normalization

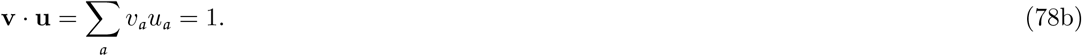

(recall that ∑_*𝒶*_ *α*_*𝒶*_ = 1). Here, we also use the following definition

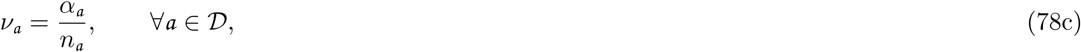

and owing to *v*_*𝒶*_ *u*_*𝒶*_ = *ν*_*𝒶*_ *n*_*𝒶*_, ∀*𝒶* ∈ 𝒟, we have

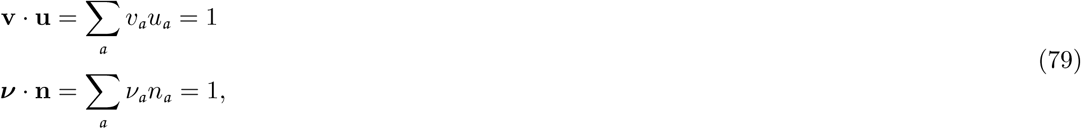

where **v, *ν*** are the vectors of *v*_*𝒶*_ and *ν*_*𝒶*_, respectively. Also, recall that *u*_*𝒶*_ *n* = *n*_*𝒶*_ and hence

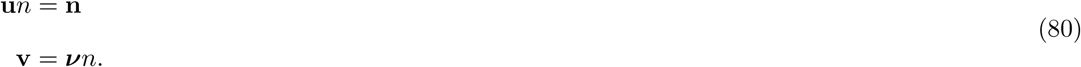

#### 6.5.1 The dynamics of *ν*_*𝒶*_

By differentiation we obtain

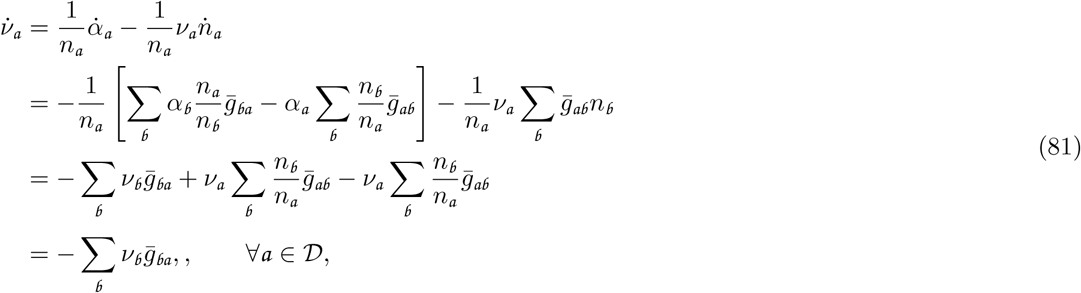

which can be expressed with a matrix notation as

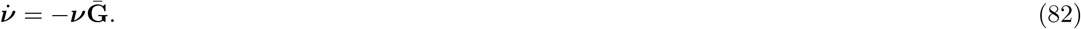

#### 6.5.2 The dynamics of *v*_*𝒶*_

Using (8), Section 6.2.1, and performing a similar calculation to above, we obtain

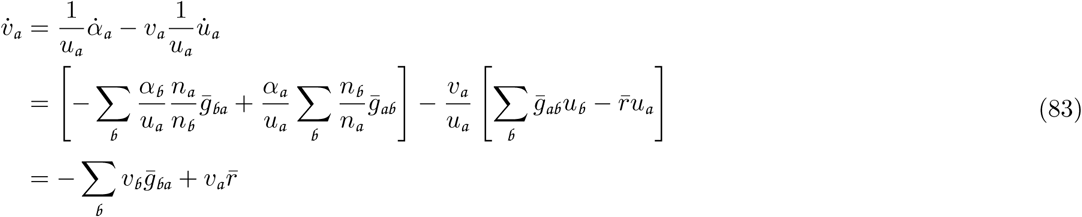

which can be expressed with a matrix notation as

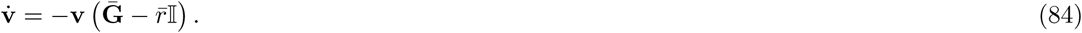

#### 6.5.3 Individual reproductive values *ν*_*𝒶*_ and *v*_*𝒶*_ as left eigenvectors

Using (82) and Section 2.4 the slow (micro-)evolutionary time definition of ***ν*** under phenotypic equality *δ* = 0 is

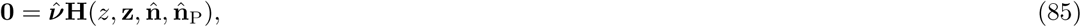

that is, 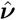 is the left eigenvector of the resident matrix 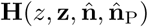 associated with the eigenvalue 0. Similarly, the slow (micro-)evolutionary time definition of **v** under phenotypic equality *δ* = 0 is

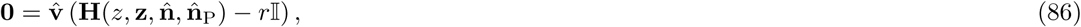

and because the identity matrix is multiplied by a scalar, the solution to above is equivalent to solving

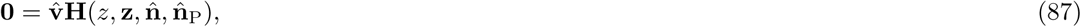

hence both 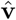 and 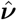 are the left eigenvectors of the resident matrix 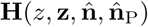 associated with the eigenvalue 0. Moreover, from (80) we have

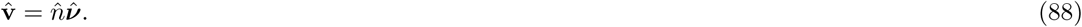

#### 6.5.4 The dynamics of the weighted mutant frequency using individual reproductive values

Here we show a more direct calculation for the dynamics of the weighted mutant frequency *p*_*α*_. Because

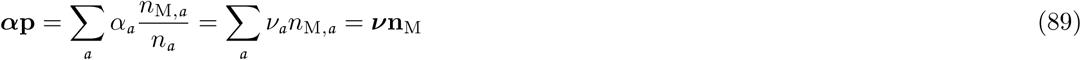

we have

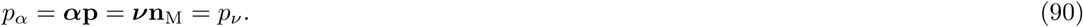

Therefore

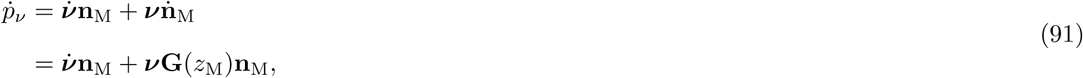

where **G**(*z*_M_) can be partitioned as

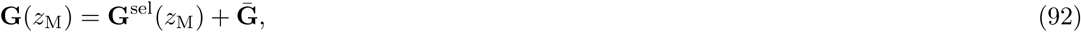

where 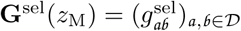 and

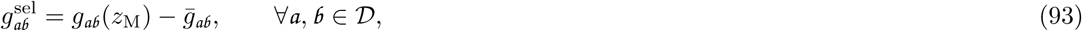

and 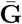 is as in the main text (5). The weighted mutant frequency can thus be written directly in terms of individual reproductive values as

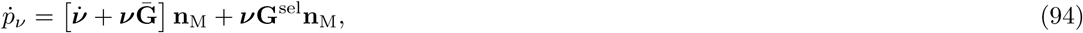

and by defining ***ν*** such that it satisfies (82) we get

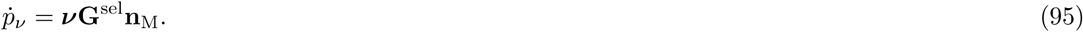

Notice that this is indeed equivalent to (18) and that under phenotypic equality 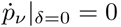. Now, taking the derivative of the above with respect to *δ* and using Section 2.4 we immediately obtain (34).

## Notes

### Competing Interest Statement

The authors have declared no competing interest.

